# Low T cell diversity is associated with poor outcome in bladder cancer: a comprehensive longitudinal analysis of the T cell receptor repertoire

**DOI:** 10.1101/2024.05.30.596555

**Authors:** Asbjørn Kjær, Nanna Kristjánsdóttir, Randi Istrup Juul, Iver Nordentoft, Karin Birkenkamp-Demtröder, Johanne Ahrenfeldt, Trine Strandgaard, Deema Radif, Darren Hodgson, Christopher Abbosh, Hugo JWL Aerts, Mads Agerbæk, Jørgen Bjerggaard Jensen, Nicolai J Birkbak, Lars Dyrskjøt

## Abstract

T cells are one of the primary effector cells in the endogenous defense against cancer, yet the clinical impact of their quantity, diversity, and dynamics remains underexplored. Here we investigated the clinical relevance of the T cell receptor (TCR) repertoire in patients with bladder cancer. In advanced-stage bladder cancer, low pre-treatment peripheral TCR diversity was associated with worse overall survival (p=0.024), particularly when it coincided with a low fraction of circulating T cells (p=0.00049). The low-diversity TCR repertoires were dominated by expanded clones that persisted throughout treatment and disproportionately targeted latent viral infections. Longitudinal analysis revealed a reduction in TCR diversity after treatment indicating an adverse effect on the immune system. In early-stage bladder cancer, we showed that immunotherapy had a stimulatory effect on TCR diversity in patients with good outcomes. Single-cell sequencing identified most expanded clones as cytotoxic T cells, while non-expanded clones were predominantly naive T cells. Overall, our findings suggest that TCR diversity is a promising new biomarker that may offer new avenues for tailored oncological treatment to enhance clinical outcomes for bladder cancer patients.

## Introduction

Cancer is generally considered a disease primarily driven by somatic alterations in the genome. However, there is now increasing evidence suggesting that cancer development and progression may also be promoted by a dysfunctional immune response^1–3^, likely shaping cancer evolution through selection for immune-resistant cancer clones. T cells are critical components of the adaptive immune system and central to the endogenous anti-tumor response. T cells carry unique T cell receptors (TCRs) generated by DNA rearrangements as they mature in the thymus. These receptors enable T cells to provide a tailored and memory-based defense by recognizing foreign antigens on the surface of antigen-presenting cells with high specificity. Proliferation following T cell activation results in the expansion of T cell clones sharing identical TCRs. This mechanism of action enables the evaluation of the T cell clone landscape through analysis of the TCR repertoire. Quantifying the TCR repertoire in circulation can reveal the breadth of a potential T cell response through analysis of TCR diversity and the strength of an ongoing response through evaluation of clonal expansion. While most TCR targets remain unknown and may be unrelated to cancer, the TCR repertoire itself can provide insights into the current state of the immune system, which may affect patient outcomes.

Bladder cancer is a highly immunogenic disease characterized by one of the highest tumor mutation burdens^4^, strong immune cell infiltration of the tumor microenvironment, and the formation of tertiary lymphoid structures within the tumor periphery. Treatment is stratified by cancer invasiveness. In high-risk non-muscle invasive bladder cancer (NMIBC), immunotherapy based on Bacillus Calmette-Guerin (BCG) instillations is the standard of care and is highly effective in preventing disease recurrence and progression^5^. Platinum-based neoadjuvant chemotherapy followed by radical cystectomy is the preferred treatment for patients with localized muscle-invasive bladder cancer (MIBC). The treatment has high perioperative morbidity and mortality, and metastatic relapse is observed in about 50% of patients^6^. For patients not eligible for chemotherapy, immunotherapy is recommended as first-line treatment^7^. Additionally, recent studies have shown that combining immunotherapy and antibody-drug conjugate (pembrolizumab and enfortumab vedotin) significantly improves outcomes compared with standard chemotherapy^8^.

As bladder cancer is highly immunogenic, investigating the TCR repertoire may significantly improve our understanding of the host anti-tumor response. Previous work has found the peripheral TCR repertoire to be associated with patient outcomes in multiple other cancer types^9–15^, indicating that the TCR repertoire affects tumor progression and may facilitate patient risk stratification. However, the impact of the peripheral TCR repertoire remains largely unexplored in bladder cancer. Additionally, the dynamics of the peripheral TCR repertoire during treatment and the interplay between tumor biology and the TCR repertoire remain underexplored.

Here, we present an in-depth analysis of the TCR repertoire in circulation to further our understanding of its biological impact and to investigate its potential for predicting clinical outcomes in patients with bladder cancer. Utilizing TCR sequencing (TCRseq), we observe that patients with low TCR diversity have significantly shorter survival. These patients often harbor large expanded T cell clones that specifically target persistent viral infections. In a single-cell analysis, we find that these large expanded T cell clones are predominantly exhausted T cells. We also find that TCR diversity is associated with distinct tumor biology, indicating that the T cell landscape may affect cancer development. Furthermore, through longitudinal analysis of TCR repertoire dynamics, we demonstrate how treatment negatively impacts both short- and long-term TCR diversity and lymphocyte abundance, particularly among patients with good outcomes.

## Results

### Patients, biological samples, and molecular data

To explore the T cell landscape, we performed TCRseq on tumor biopsies and longitudinal blood samples from patients with MIBC (n = 119) and NMIBC (n = 30). This was analyzed together with whole exome (WES), whole genome (WGS), and transcriptome (RNAseq) sequencing data on patient subsets available from previous analyses^1,16–18^ (**Figure 1A**, data overview in **Figure S1**). Patients were treated and followed according to standard clinical guidelines at Aarhus University Hospital, Denmark. A summary of clinical and histopathological characteristics is provided in **Table S1**.

**Fig. 1.**
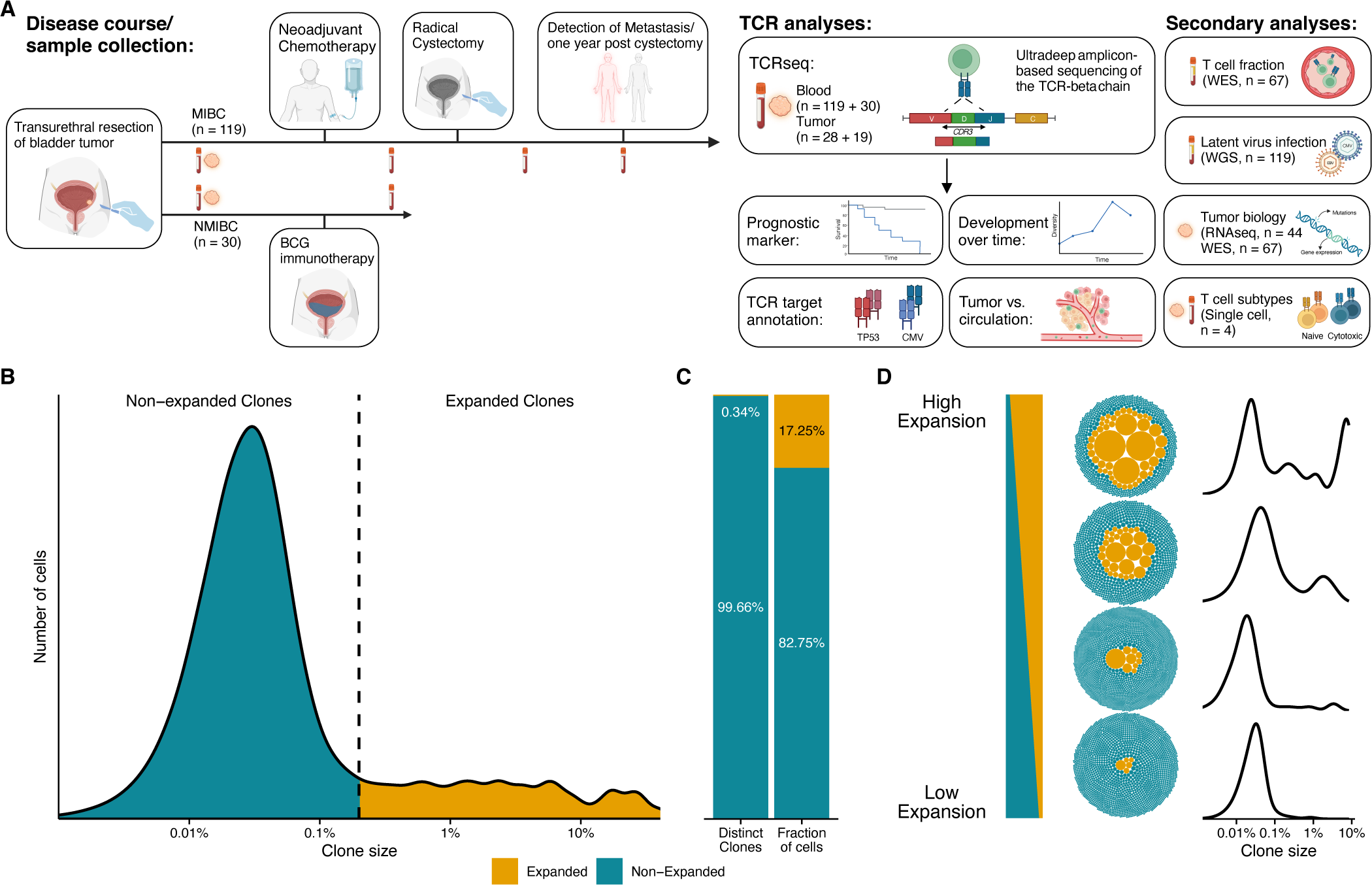
Study overview and baseline TCR landscape. **A**, Overview of the patient cohorts and study analyses. Patients with MIBC (n = 119) and NMIBC (n = 30) were included for peripheral TCRseq. A subset of patients were subject to longitudinal (MIBC, n = 33; NMIBC, n = 28) and tumor TCRseq (MIBC, n = 28; NMIBC, n = 19). Blood vials and tumors indicate the time points for the collection of blood and tumor samples, respectively. Created with BioRender.com. **B**, Clone size distribution in blood at baseline for patients with MIBC (n samples = 119, n clones = 635,814). Clones representing more than 0.2% of cells in a repertoire were defined as expanded. The density plot is weighted by clone size to represent the distribution of cells. **C**, The number of expanded clones among all clones and the total size of expanded clones among all T cells (representing the overall relative size of expanded clones). All repertoires were concatenated for this analysis. **D**, Visualization of varying levels of clonal expansion using bubble plots of representative repertoires (patients 82, 8, 102, and 11). The T cell repertoire is represented as a collection of bubbles, and the size of a bubble represents clone size. Colored according to expansion threshold. Density plots: as described in B, but each for a repertoire from a single patient. (See also **Figures S1, S2** and **Table S1**).

### TCR clonal expansion varied substantially among patients with MIBC prior to chemotherapy

We investigated the peripheral blood TCR landscape before chemotherapy (baseline) in 119 patients with MIBC using TCRseq on buffy coat DNA. This resulted in a median recovery of 4908 unique TCR CDR3β chains (hereafter referred to as T cell clones) per sample (range 1479-15,679). Analysis of the combined clone size distribution across all patients revealed a right-skewed distribution composed of two distinct categories of T cell clones: small non-expanded T cell clones, each clone represented by few T cells, and large expanded T cell clones, each clone represented by many T cells (**Figure 1B**). Using a previously defined threshold^19^, we categorized the T cell population into expanded clones, represented at a frequency exceeding 0.2%, and non-expanded clones, represented at a frequency below 0.2%. The group of expanded clones constituted only 0.34% of the total amount of unique T cell clones while representing 17.25% of the total fraction of T cells (**Figure 1C**). We observed a considerable variation in clonal expansion across patients (range 2%-65%), corresponding to individual repertoires ranging from almost no clonal expansion to repertoires dominated by expanded T cell clones (**Figures 1D, S2**).

### Low TCR diversity and low T cell fraction at baseline are associated with worse disease outcomes in MIBC

To assess the clone-size distribution without a specific threshold for clonal expansion we utilized the normalized Shannon diversity index as a measure of TCR diversity, which strongly correlated with the fraction of expanded T cell clones (r = -0.98, p < 0.0001; **Figure S3A**). Baseline TCR diversity was significantly associated with survival. Patients with below median TCR diversity experienced significantly shorter overall survival (OS; HR = 2.3, p = 0.024; **Figure 2A**). They also had shorter recurrence-free survival (RFS), although it did not reach statistical significance (HR = 1.7, p = 0.22; **Figure S3B**). Patients who developed metastatic disease had significantly lower TCR diversity compared with those who did not develop metastasis (p = 0.036; **Figure S3C**), although no association was found between baseline TCR diversity and chemotherapy efficacy (pathologic downstaging and circulating tumor DNA (ctDNA) clearance; **Figures S3D,E**).

**Fig. 2.**
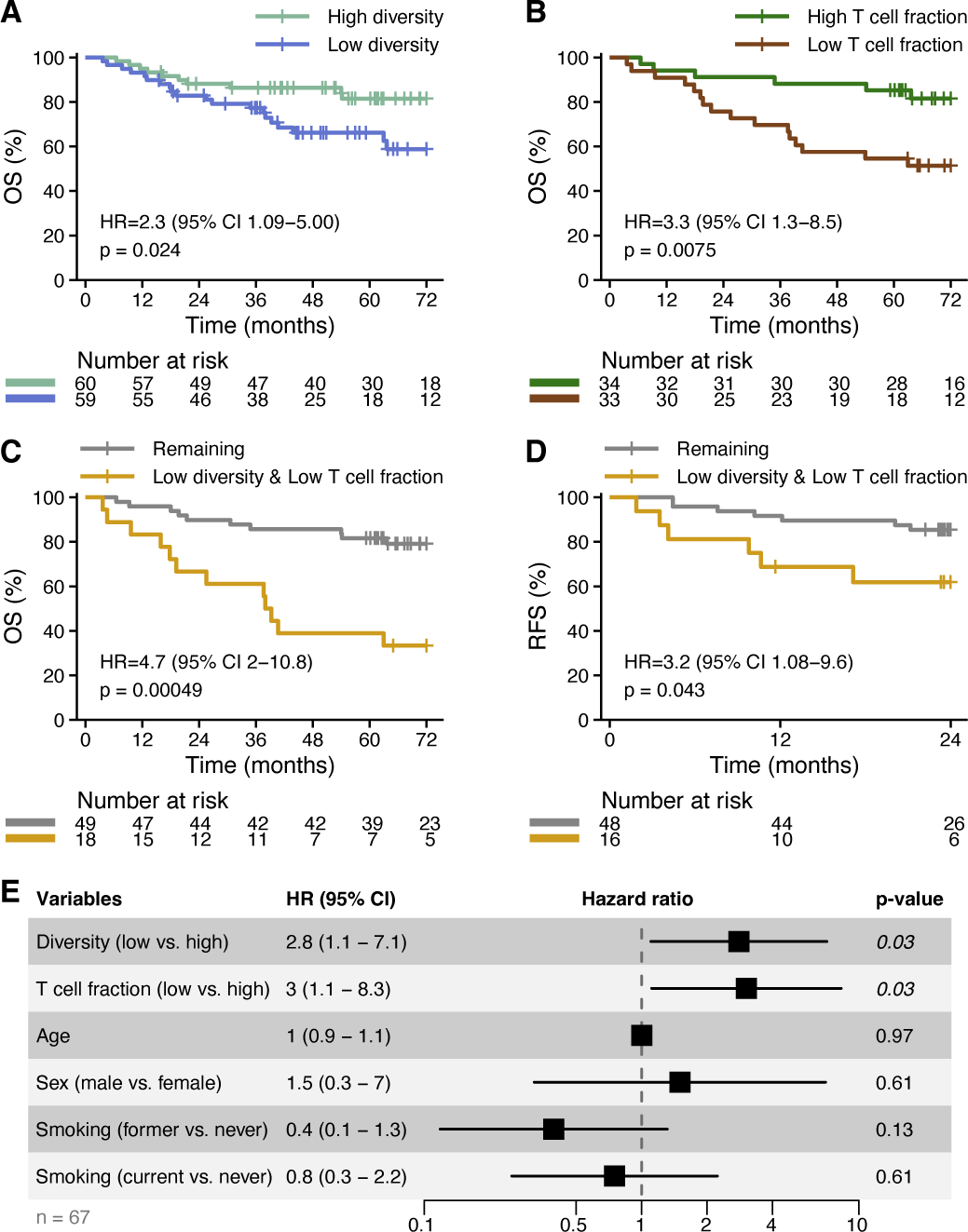
Baseline TCR diversity and T cell fraction are associated with overall survival. **A**, Survival analysis associating TCR diversity (median split) with OS. **B**, Survival analysis associating T cell fraction (median split) with OS. **C**, Survival analysis of OS comparing patients with low normalized Shannon diversity index (below median) and low relative T cell fraction (below median) to all other patients. **D,** Survival analysis of RFS comparing patients with low normalized Shannon diversity index (below median) and low relative T cell fraction (below median) to all other patients. **E,** Forest plot of multivariable analysis including TCR diversity, T cell fraction, age, sex, and smoking status. CI: confidence interval. (See also **Figures S3, S4**).

TCR diversity measures the relative distribution of T cell clones but contains no information about the amount of T cells in circulation. To investigate the associations between TCR diversity, T cell levels, and outcomes, we determined the relative fraction of T cells in circulation before treatment in 67 patients with MIBC using TcellExTRECT^20^ on germline buffy coat WES data. Patients with a below-median T cell fraction had a significantly shorter OS (HR = 3.3, p = 0.0075; **Figure 2B**) and RFS (HR = 4.2, p = 0.016; **Figure S3F**). As shown for TCR diversity, T cell fraction was not associated with chemotherapy efficacy (**Figures S3G,H**), yet patients who developed metastasis had a significantly lower fraction of T cells compared to patients who did not (p = 0.022; **Figure S3I**).

Interestingly, TCR diversity and T cell fraction were uncorrelated (Spearman’s rho = 0.032, p = 0.8; **Figure S3J**), indicating that they represent independent measures of the T cell repertoire. Neither measure was associated with patient characteristics (**Figures S3K-T**), although most patients in the lowest TCR diversity quartile were older than 60 years (28/30 patients, p = 0.02; **Figure S3O**). In a combined analysis, we found that patients with both low TCR diversity and low T cell fraction had shorter OS (HR = 4.7, p = 0.00049; **Figures 2C, S3U**) and shorter RFS (HR = 3.2, p = 0.043; **Figures 2D, S3V**) compared to the remaining patients. TCR diversity and relative T cell fraction were identified as the only significant predictors of OS in both univariate and multivariable analyses **(Figures 2E, S3W**). Together, these results indicate that peripheral blood TCR diversity and T cell fraction are both independently associated with outcomes for patients with MIBC and that the combination of low TCR diversity and low T cell fraction represents a particularly poor prognosis.

Lastly, we aimed to validate these findings in an independent cohort of patients with non-metastatic MIBC (stage I-III patients from TCGA cohort)^21^. As deep TCRseq data were unavailable, we used an orthogonal approach to determine TCR diversity. We utilized germline WES data to determine both peripheral TCR diversity and T cell fraction. Of 262 samples, 107 had sufficient TCR sequences to estimate TCR diversity. We observed that low TCR diversity and low T cell fraction were associated with shorter OS both combined and individually, analogous to the main cohort (p = 0.01, p = 0.017, p = 0.067; **Figures S4A-C**). In the same manner, we determined TCR diversity and T cell fraction in a cohort of patients treated with immunotherapy for metastatic bladder cancer (IMvigor210 cohort)^22^. We again observed that patients with both low TCR diversity and low T cell fraction had significantly shorter OS (HR = 3.0, p = 0.042; **Figure S4D**), although here neither low TCR diversity nor low T cell fraction was associated with outcome individually (**Figures S4E,F)**

### Clonal hematopoiesis of indeterminate potential (CHIP) does not induce low TCR diversity

To investigate if the observed T cell clonal expansion might be induced by lymphoid CHIP, we determined the prevalence of CHIP-associated somatic mutations in the patients using germline WES data. We defined CHIP as previously described^23^, requiring at least one CHIP-associated mutation observed at a minimum frequency of 2%. We detected CHIP in 13% (9/67) of patients with MIBC. These patients had significantly higher baseline TCR diversity than those without CHIP (p = 0.016; **Figure S5A**), indicating that CHIP does not promote clonal expansion of T cells. CHIP was neither associated with T cell fraction nor clinical outcomes (**Figures S5B-D**).

### Expanded T cell clones primarily target antigens from latent viral infections

Next, we explored the antigen targets of the T cell clones from the baseline MIBC samples. First, we established clusters of highly homologous TCR sequences based on sequence similarity, using GLIPH2^24^. Likely antigen targets of these clusters were then inferred based on sequence similarity with known CDR3β-antigen pairs^25^ (**Figure 3A**). We found that the TCRs of expanded clones were more likely to have an inferred target relative to TCRs of non-expanded clones (odds ratio (OR) = 2.55, p = 3x10^-8^; **Figure 3B**). We observed a marked difference between the inferred antigen targets of expanded and non-expanded clones (**Figure 3C**), with expanded clones significantly more likely to target Epstein-Barr virus (EBV; adjusted p < 0.0001) and cytomegalovirus (CMV; adjusted p = 0.0007) relative to non-expanded clones (**Figure 3D**). Many viral infections, including CMV and EBV, may persist as latent infections inside cells after the primary infection has been resolved^26^. To investigate if latent viral infections drove the T cell clonal expansion, we explored the association between TCR diversity and EBV and CMV. As the serostatus for CMV and EBV of the patients were unknown, we constructed a pipeline utilizing Kraken2^27^ to detect viral DNA based on WGS data from cell-free DNA in plasma samples. We found DNA evidence of persistent EBV and CMV infections in 43% (51/119) and 54% (64/119) of patients with MIBC, respectively. This matches the reported seroprevalence of CMV (58%)^28^, but underestimates that of EBV (95%)^28^. Interestingly, patients with detectable CMV DNA had significantly lower TCR diversity (p = 5.3x10^-5^; **Figure 3E**) while detectable EBV DNA showed no association (**Figure S5E**). Patients with detectable CMV infection had a higher fraction of their TCRs targeting CMV antigens than those with no detectable CMV (**Figure S5F**). Latent CMV infection was neither associated with T cell fraction nor disease outcome (**Figures 3F, S5G,H**). In a linear regression model predicting TCR diversity, both CMV and metastatic disease were significant variables, suggesting that both factors impact TCR diversity independently (**Figure S5I**). These results indicate that the majority of expanded T cell clones target pathogens, and are thus likely to be non-cancer specific.

**Fig. 3.**
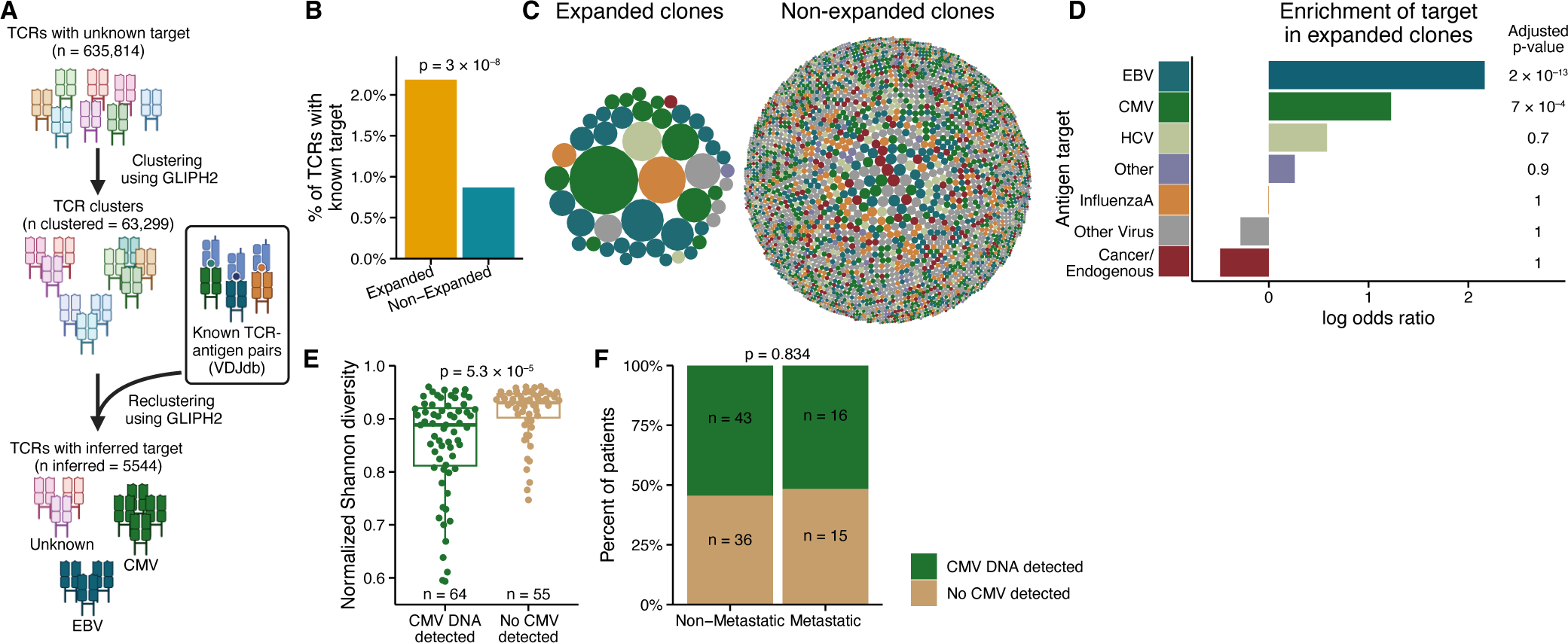
Expanded TCR clones disproportionately target viral antigens. **A-D,** TCR target inference by TCR sequence-homology clustering using GLIPH2. **A**, Overview of the annotation pipeline. First, baseline MIBC TCR clones were clustered based on sequence homology using GLIPH2. The antigen targets of these clusters were then inferred by reclustering all TCRs together with TCR sequences from known TCR-antigen pairs obtained from VDJdb. Only clusters reemerging when reclustered were further considered. TCR clones in clusters encompassing a TCR with a known antigen were assumed to target the same antigen. Created with BioRender.com. **B**, Percentage of expanded and non-expanded TCR clones that are included in TCR clusters together with a TCR sequence with a known antigen target. Test for association between inclusion in a cluster with an inferred target and a clone being expanded. All clones concatenated for analysis (non-expanded, n = 633,663; expanded, n = 2151). **C**, Visualization of inferred targets for expanded and non-expanded clones. Only TCRs with an inferred target are visualized. **D**, Test for target enrichment in expanded clones, based on comparing the amount of expanded and non-expanded clones with a given target relative to the total amount of expanded and non-expanded clones (non-expanded, n = 633,663; expanded, n = 2151). **E-F**, Detection of DNA from CMV in plasma samples, analyzed using Kraken2 on WGS data from patients with MIBC. **E**, Association between normalized Shannon diversity index and CMV DNA detection. The center line is the median, box limits represent upper and lower quartiles, and whiskers represent 1.5 times the interquartile range. **F**, Association between the development of metastatic disease and CMV DNA detection. (See also **Figure S5**).

### Expanded clones are highly persistent throughout the disease course

To explore the TCR repertoire during the disease course, we analyzed longitudinal blood samples from 33 patients with MIBC. Additional TCRseq was performed on samples taken after chemotherapy, three weeks after cystectomy, and either at metastatic relapse or one year after cystectomy (3-4 samples per patient). Clones were defined as persistent if found in all available samples, recurrent if found in more than one sample, and transient if found in only one sample. We observed considerable variation across patients at baseline (**Figure 4A**) and a sharp increase in the prevalence of persistent clones with increasing clone size (**Figure 4B**). The majority of expanded clones were expectably categorized as persistent while non-expanded clones were commonly transient (**Figures 4C, S6**). Thus, within the study timeframe (median 16 months, range 4-64), almost all expanded T cell clones detected at baseline remained detectable in circulation. This suggests that the overall T cell landscape remains relatively stable, and significant contractions of expanded T cell clones are infrequent or occur slowly. The amount of these clones was clinically relevant, as the patients who developed metastatic disease had a higher fraction of persistent and recurrent clones (**Figure 4D**). Consistent with this, we found that patients with high amounts of persistent and recurrent clones had shorter OS (**Figure S7A**).

**Fig. 4.**
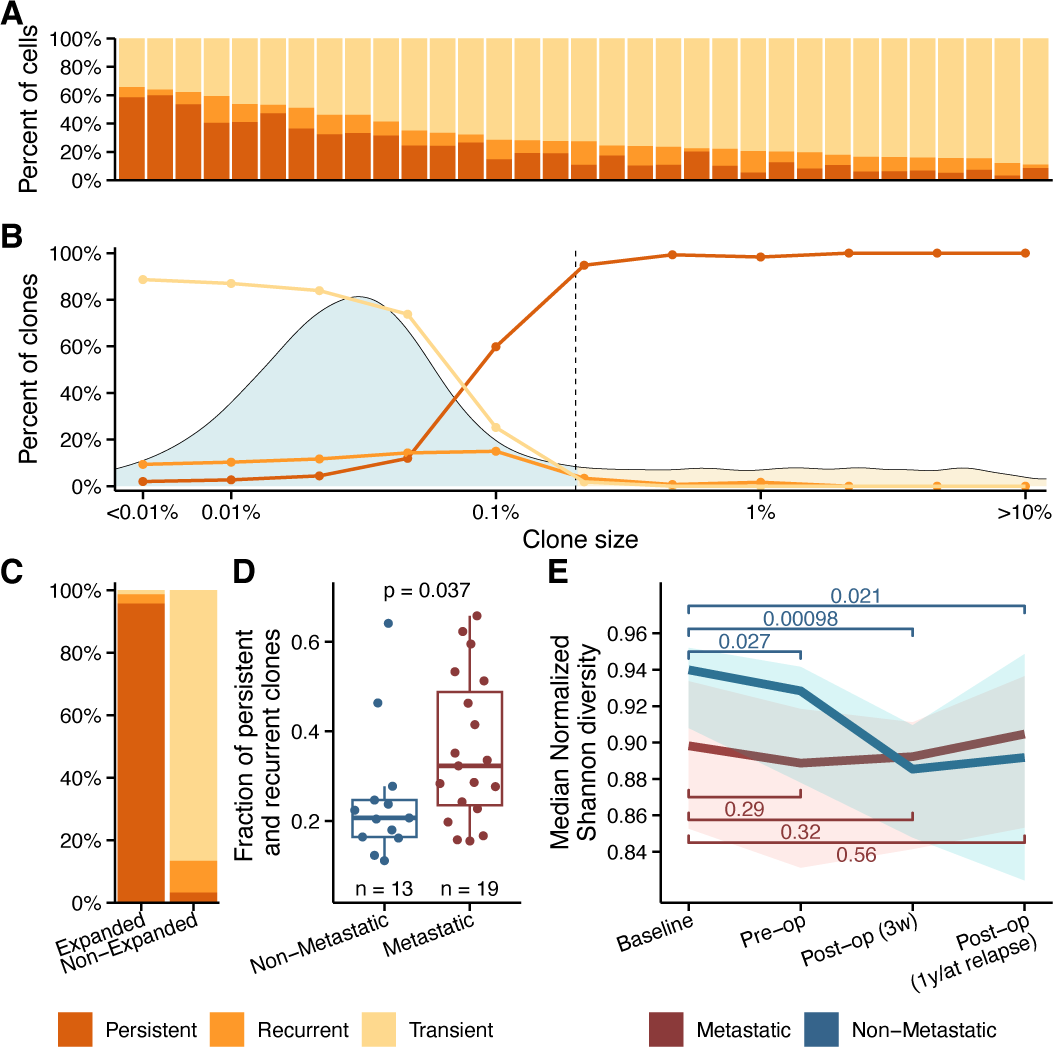
Longitudinal analysis of the TCR landscape. Analysis of the TCR repertoire through treatment for a subset of patients with MIBC (n = 33). Clones were defined as persistent if found at all available time points, recurrent if found at more than one time point, and transient if only found once. All patients had 3-4 samples available: 1) Baseline, taken before treatment initiation, 2) Pre-op, taken between chemotherapy and cystectomy, 3) Post-op (3w), taken three weeks after cystectomy, and 4) Post-op (1y / at relapse), taken either at detection of metastasis or one year after cystectomy for patients that with and without metastasis, respectively. **A,** Visualization of the abundance of persistent, recurrent, and transient clones at baseline. Each bar corresponds to one patient. **B,** Association between clonal persistence and size of clones. Lines indicate the percentage of clones in each category at a specific size bin. Bins are uniformly distributed on log scale, x-axis placement indicates the end of the bin. All clones were concatenated for this analysis. The density plot shows the baseline size distribution of T cell clones (from **Figure 1B**). The dashed line as well as the blue and yellow colors indicate the threshold for defined expanded T cell clones. **C**, The percentage of expanded and non-expanded clones divided into persistent, recurrent, and transient clones. Expanded clones are primarily persistent, while non-expanded are primarily transient. **D**, Box-plot showing the association between the development of metastatic disease and the total fraction of persistent and recurrent clones at baseline (representing the relative size of persistent and recurrent clones; one patient excluded from analysis due to incomplete follow-up). **E**, Lineplot showing the change in median normalized Shannon diversity index through treatment for patients with and without metastatic disease. The shadow behind the lines shows the interquartile range. P-values are calculated using the individual normalized Shannon diversity index values. (See also **Figures S6, S7**).

### Treatment decreases TCR diversity and lymphocyte counts in patients with good outcome

We analyzed the dynamic changes to the TCR landscape during and after treatment in patients with metastatic disease and those without, separately. While patients in the non-metastatic group initially exhibited higher TCR diversity, these patients experienced a decrease in diversity during treatment. This was not observed in the group of patients with metastatic disease, resulting in equivalent TCR diversity in both patient groups after cystectomy (**Figures 4E, S7B**).

To further investigate the dynamics of the immune landscape, we analyzed longitudinal biochemical laboratory measures of blood cell counts from 58 patients with MIBC. We found that lymphocyte counts, which are primarily T cells, consistently decreased from baseline throughout treatment in patients without metastatic disease, a trend not evident in the patients with metastatic disease (**Figure S7B**). Contrary, neutrophil counts and overall leukocyte counts only decreased after chemotherapy initiation, whereafter they recovered to baseline levels in both patient groups (**Figures S7D,E**). These results indicate that treatment may negatively impact TCR diversity and lymphocyte counts, particularly in good prognosis patients with high-diversity repertoires.

### TCR diversity is associated with outcomes in early-stage bladder cancer

To investigate the impact of TCR diversity on early-stage bladder cancer, we performed TCRseq on blood samples taken before and after BCG immunotherapy from 30 patients with NMIBC (**Figures 1A, S8**). Noticeably, the TCR diversity was equivalent to that of patients with MIBC (**Figure S9A**). Outcomes after treatment were dichotomized into either late or no high-grade recurrence (> two years), or early high-grade recurrence (< two years) or progression. We observed no significant difference in TCR diversity between the two groups, before or after BCG (**Figure S9B**). Although we did not find an overall change in diversity after BCG (**Figure S9C**), we found that the TCR diversity increased significantly in patients with late or no high-grade recurrence, contrasting the chemotherapy-treated MIBC cohort where a decrease in diversity was observed (**Figure S9D**). Progression-free survival (PFS) was not associated with TCR diversity before BCG (median split; **Figure S9E**). However, patients with low diversity after BCG had shorter PFS (HR = inf, p = 0.015; **Figure S9F**). We estimated the T cell fraction for 110 patients with NMIBC from an extended cohort utilizing WES data applicable for TcellExTRECT and found that patients with a lower T cell fraction had shorter PFS (HR = 3.8, p = 0.027; **Figure S9G**), which is comparable to the MIBC cohort. These findings collectively support that low TCR diversity and a low T cell fraction indicate a poor prognosis in NMIBC.

### Peripheral TCR diversity affects tumor biology

To evaluate if peripheral TCR diversity was associated with specific tumor biology characteristics, we investigated RNAseq and WES data from tumor biopsies from patients with MIBC. We performed differential gene expression analysis on the 2000 most variable genes. Of these, we found that 153 and 39 genes were significantly upregulated in the patients with low (n = 23) and high (n = 21) TCR diversity, respectively (**Figure 5A**). Genes upregulated in patients with low TCR diversity were mainly associated with extracellular matrix organization or signal transduction. Genes upregulated in patients with high TCR diversity were related to the metabolism of proteins or RNA (**Figures 5B, S10A**). When investigating tumor genetics, we found no association between peripheral TCR diversity and the frequency of common bladder cancer driver genes nor the overall tumor mutation burden (**Figures S10B,C**). Lastly, we explored if TCR diversity might affect the ability to detect ctDNA in baseline blood samples. Pre-treatment ctDNA has previously been associated with aggressive disease and increased risk of metastatic progression^16^. Interestingly, we found that TCR diversity was significantly lower in the ctDNA-positive group, supporting that patients with low TCR diversity may harbor tumors with an increased risk of metastatic dissemination (**Figures S10D**).

**Fig. 5.**
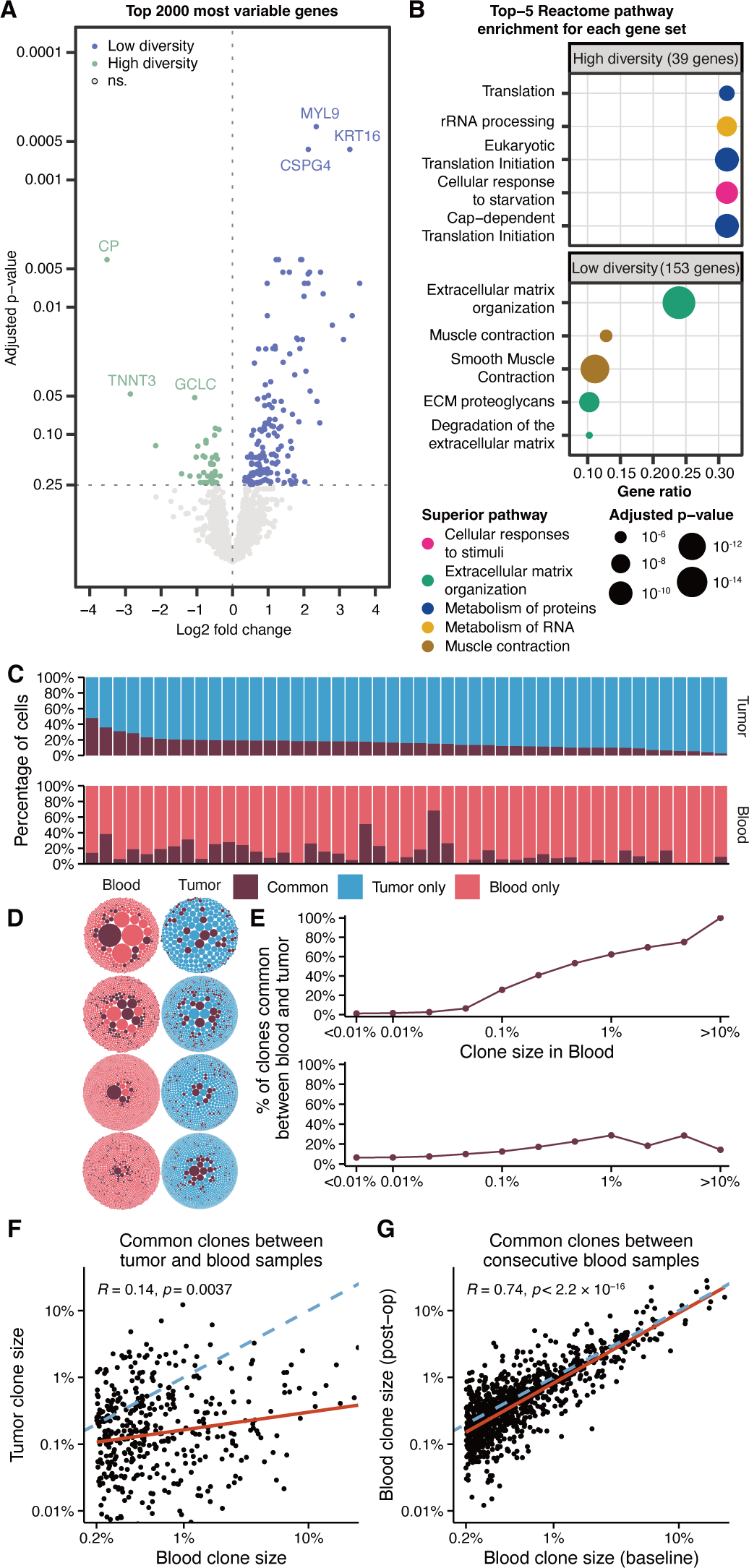
Effect of peripheral TCR diversity on MIBC tumor biology and exploration of tumor TCR repertoires. **A-B**, Analysis of tumor biology in relation to peripheral TCR diversity for patients with MIBC. **A**, Differential gene expression analysis of top 2000 most variable genes comparing patients with high and low TCR diversity. **B**, Reactome pathway enrichment analysis using the significantly differentially expressed genes with more expression in patients with high TCR diversity (n = 39) and low TCR diversity (n = 153) as input. The plot shows the top five pathways for each set of genes. **C-E**, Joint analysis of paired tumor and blood repertoires for MIBC (n = 28) and NMIBC (n = 19). **C**, The amount of overlap between blood and tumor TCR repertoires (representing the relative size of common clones). Vertically aligned bars represent one patient. **D**, Visualization of clones common between blood and tumor repertoires. Horizontally aligned bubble plots represent the same patient (patients 82, 8, 102, and 11). **E**, The percentage of clones that are common between blood and tumor at varying clone sizes in either blood (top) or tumor (bottom). As clones become larger in the blood they are more likely to also be found in the tumor. This is less likely for larger clones in the tumor. Each point represents a uniformly distributed bin (log scale), x-axis placement indicates the end of the bin. All common clones across patients were concatenated for this analysis. **F-G,** Correlation of size between expanded clones at baseline shared with either tumor (**F**, n = 423) or post-op blood (**G**, n = 973) (post-op includes after BCG for NMIBC and three weeks after cystectomy for MIBC, only including persistent clones). The blue dashed line indicates a one-to-one linear relationship, while the orange line indicates a linear model fit. (See also **Figures S10, S11**).

### Tumor TCR repertoires are distinct from peripheral repertoires

The relationship between peripheral blood and tumor TCR repertoires was investigated by performing additional TCRseq on tumor DNA from 47 patients (MIBC = 28, NMIBC = 19). Tumor TCR repertoires generally had fewer clones and were less diverse than the peripheral blood samples (**Figures S10E,F**). Tumor TCR diversity was not associated with outcome in either cohort (**Figures S10G,H**). Neither the number of clones nor TCR diversity were correlated between tumor and peripheral blood TCR repertoires, indicating that the two repertoires represent different biologies (**Figures S10I,J**). To explore this further, we analyzed the clones that were common between the baseline blood repertoires and the tumor repertoires (**Figure 5C**). Visualization of the common clones revealed that tumor and blood shared limited amounts of T cell clones (**Figures 5D, S11**). However, we noticed that larger clones in the blood had an increased tendency to be common, while this was less pronounced for larger clones in the tumor (**Figure 5E**). To investigate if T cell clones expanded in the blood were targeting the tumor microenvironment, we compared the sizes of the T cell clones common between blood and tumor. The size of the common clones exhibited a weak correlation between tumor and blood (**Figure 5F**), and a strong correlation between consecutive blood samples (**Figure 5G**). Common clones were generally smaller in the tumor than in the blood, indicating that expanded clones infiltrate the tumor less than expected at random, presumably because other T cell clones preferably infiltrate the tumor.

### Single-cell sequencing reveals the T cell subtypes of expanded and non-expanded clones

To gain information on the T cell subtypes in patients with bladder cancer we performed single-cell RNAseq on paired tumor and blood samples. Given the requirement for viable cells, we collected fresh blood and tumor samples from four patients with bladder cancer undergoing cystectomy. Paired TCR and full-length RNA profiling were performed on isolated T cells resulting in cell recovery ranging between 271-3199 cells, and clone count varying between 226-2248 clones (**Figure S12A**). When analyzing T cell clones common between blood and tumor samples, we saw similar patterns as in the bulk data. This supports that the chance for a clone to be common largely depended on its size in peripheral blood (**Figures 6A,B**).

**Fig. 6.**
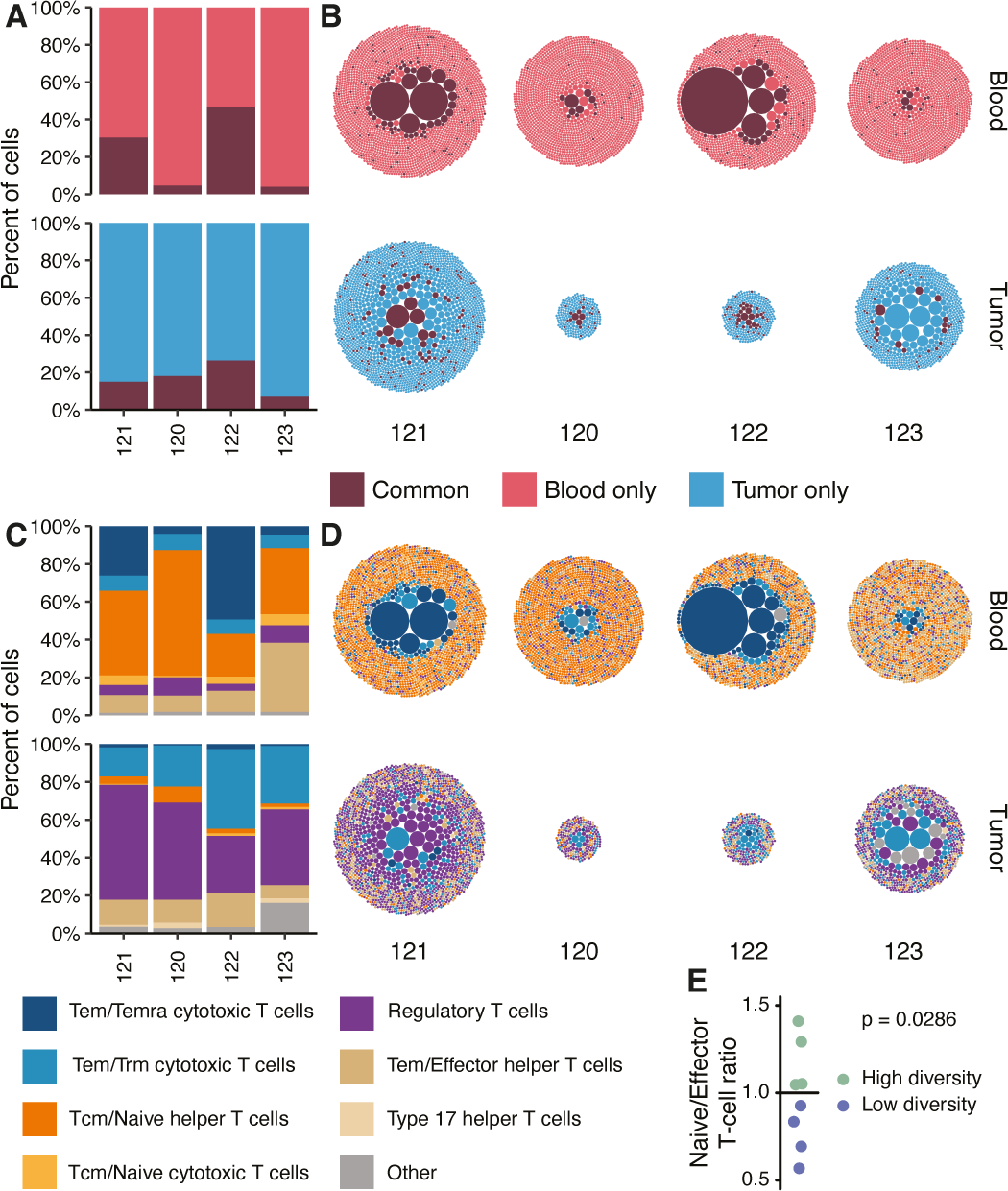
Inference of T cell subtypes of expanded and non-expanded clones. Single-cell analysis of paired tumor and blood samples from four patients with bladder cancer. **A-D,** Each patient is shown in vertically aligned bars/bubbles with blood samples on top and tumor samples below. **A,** Relative proportion of clones common between tumor and blood, and clones unique to tumor or blood. **B,** Visualization of the common clones. **C,** Relative proportion of eight different T cell subtypes in each sample. **D,** Visualization of the T cell subtypes. Other include Trm cytotoxic T cells, Memory CD+ cytotoxic T cells, Treg(diff), Follicular helper T cells, Type 1 helper T cells, CRTAM+ gamma-delta T cells, gamma-delta T cells, MAIT cells, Cycling T cells, CD8a/b(entry), and Double-positive thymocytes. Tem: effector memory T cell. Temra: effector memory T cell reexpressing CD45RA. Trm: tissue-resident memory T cell. Tcm: central memory T cell. Treg(diff): differentiating regulatory T cell. MAIT: mucosal-associated invariant T cell. CD8a/b(entry): developing CD8 alpha-beta T cell (late double-positive stage). **E,** Ratio of naive/effector T cells for eight patients with RNAseq from blood showing higher ratios in patients with high TCR diversity relative to patients with low TCR diversity. (See also **Figure S12**).

The T cell clones were annotated using CellTypist^29^ to identify T cell subtypes revealing a clear contrast between the blood and tumor samples. Regulatory T cells were dominant in the tumor samples, while a combination of naive and cytotoxic T cells were dominant in the blood samples (**Figure 6C**). More explicitly, the majority of the expanded clones found in the blood samples were annotated as terminally differentiated effector memory (Temra) T cells with high levels of cytotoxic and exhausted genes whereas the non-expanded clones were largely naive T cells (**Figures 6D, S12B**). Cytotoxic T cells were most likely to be common between two samples, both in tumor and blood (**Figure S12C**).

To examine if the expanded T cell clones found by bulk TCRseq analysis also matched a profile of cytotoxic T cells, we performed RNAseq on buffy coat samples from eight patients with MIBC, four with high and four with low TCR diversity as measured by bulk TCRseq. We estimated the immune cell composition of each patient through Gene Set Variation Analysis utilizing signatures from Travaglini et al^30^. This revealed a lower naive to cytotoxic T cell ratio in patients with low TCR diversity relative to patients with high TCR diversity (**Figure 6E**). This supports that the majority of expanded clones found in patients with low TCR diversity may primarily be composed of cytotoxic T cells, as observed in the single-cell data.

## Discussion

In this study, we present a comprehensive analysis of the TCR repertoire in patients with bladder cancer. We have characterized the peripheral and tumor TCR landscapes and explored the dynamics of peripheral TCR repertoires during treatment. Our analysis of the peripheral blood T cell landscape revealed that both the number of T cells and the diversity of TCR landscapes are important for the survival of patients with bladder cancer. We have specifically shown that both low TCR diversity and low relative T cell fraction are associated with poor patient outcomes in MIBC and NMIBC. These findings were validated in independent cohorts from TCGA and IMvigor210. Although, for these cohorts TCR diversity was estimated using germline WES data as TCRseq data was not available, resulting in lower recovery of T cell clones. To the best of our knowledge, we are the first to demonstrate a favorable association between high TCR diversity in peripheral blood and the outcome of bladder cancer and to show that combining TCR diversity and relative T cell fraction improves patient stratification. High peripheral blood TCR diversity has previously been reported to be associated with improved outcomes in other cancer types, including melanoma^9^, renal^10^, cervical^11^, lung^12,13^, and breast cancer^14,15^. Together this underlines that TCR diversity is likely a generalizable measure of the state of the immune system, which may impact cancer outcomes.

In our study, expanded T cell clones were inferred to disproportionately target EBV and CMV antigens. CMV has previously been reported to promote the deterioration of T cell immunity by the formation of expanded T cell clones^31^, and we showed that detection of CMV DNA was associated with decreased diversity. While CMV detection was not associated with outcomes in our cohort, CMV has previously been linked to increased mortality^32^. The absence of a discernible effect of EBV could be caused by reduced sensitivity to detect EBV infections. Although EBV has been associated with cancer formation^33^, almost all individuals have previously been infected with EBV^28^ making quantification of its effect challenging in the present study. Further studies of viral infections and their impact on immune health and oncological treatment regimes are urgently needed.

Based on longitudinal analysis, we observed that the patients with MIBC who did not develop metastasis showed a significant decline in peripheral blood TCR diversity and lymphocyte counts. These measures did not fully recover during the study, indicating that these patients may have suffered permanent treatment-induced immune degradation. In contrast, no systematic changes were observed in patients developing metastatic disease. This suggests that while patients with a healthier immune system are less likely to develop metastases, their immune systems are more affected by treatment. These effects may be permanent, and could adversely affect patient health in the years to come, potentially leading to an increased risk of severe infections and risks of developing other cancers. The reduced TCR diversity and lymphocyte count are likely caused by cytotoxic cisplatin chemotherapy. While highly effective against cancer cells, cisplatin may cause degradation of the thymus^34^, causing reduced production of naive T cells with potentially detrimental effects on long-term patient health^35^. Conversely, for patients with early-stage bladder cancer, we found that patients with good outcomes after BCG showed an increase in TCR diversity. This increase may stem from the non-tumor-specific immune stimulation by BCG, reflecting an important functional aspect of BCG response. Taken together, this work suggests that a level of restraint should be considered when administering chemotherapy. Particularly, future clinical trials should investigate if increased surveillance, e.g. using ctDNA to identify early relapse^16^, might be a suitable alternative for patients with an otherwise good prognosis and a healthy immune system. While we find an association between low TCR diversity and poor outcomes in bladder cancer, our study has limited power to determine the cause/effect relationship between TCR diversity and disease progression. However, we observed no difference in TCR diversity between different T and N stages in MIBC, nor between MIBC and NMIBC. Together with the lack of systematic changes in TCR diversity over time in patients developing metastatic disease, this indicates that disease progression has little impact on TCR diversity.

TCR diversity in circulation also affected tumor biology. We showed that genes with higher expression in patients with low TCR diversity were mainly related to extracellular matrix organization. Remodeling of the extracellular matrix is known to be related to cancer progression and metastatic disease^36^, which supports our finding that low TCR diversity is associated with poor outcomes. In addition, we showed that the TCR repertoire found in circulation was distinct from the repertoire found within the tumor. The limited amount of common clones between tumor and blood may be caused by our TCRseq approach, which utilized buffy coat samples without enriching for tumor-targeting T cells. These cells are known to be present in minute amounts in peripheral blood^37^. Using single-cell analysis, we showed that clones common between tumor and blood were mainly cytotoxic T cells, consistent with previous work in four patients with renal and lung cancers^38^. Furthermore, we demonstrated that the expanded clones found in circulation were dominated by cytotoxic T cells, mostly with an exhausted phenotype. Conversely, non-expanded clones were mainly naive T cells, which could indicate higher thymic activity with ongoing production of new T cells in patients with high TCR diversity. The main components in the tumor T cell landscape were regulatory T cells. These produce an immunosuppressive tumor microenvironment, likely indicative of the tumor having escaped immune surveillance. Our single-cell data analysis further demonstrated a large difference in T cell composition between peripheral blood and tumor tissue. Given the link between the circulating TCR repertoire and prognosis, this suggests that general immune health can be assessed based on circulating T cells from a minimally invasive blood sample, potentially reflecting the patient’s overall immune capacity.

Collectively, we provide evidence that the T cell repertoire in circulation is highly diverse among patients, and is associated with outcome - indicating a potential impact on cancer development. Considering the general health relevance of a well-functioning immune system, these findings may have significant clinical implications on risk stratification and treatment sequencing. Immune-competent patients with high TCR diversity might benefit from immune-boosting therapies such as immune checkpoint inhibitors, improving their outcomes even further. In addition, chemotherapy strategies should potentially be used primarily in patients with low TCR diversity and hence low immune competence. Indeed, TCR diversity and other methods to assess patient immune competency may be highly predictive of immunotherapy response, given that a competent immune system is likely a prerequisite for mounting an anti-cancer response. Thus, developing accurate measures to assess immune competency may significantly improve patient stratification for anticancer therapies across various cancer types. Future clinical trials should focus on integrating immune competency measures into precision medicine approaches to optimize anti-cancer therapies and patient survival.

## Materials and Methods

### Human participants

National Scientific Ethical Committee granted permission to perform this project (#1706291; #1302183; #1708266). Written informed consent was obtained from all patients before inclusion in the study. The study’s main cohort consists of 119 patients diagnosed with localized MIBC, prospectively enrolled between 2013 and 2022. Blood samples were collected at uniformly scheduled clinical visits during a two-year follow-up period. As part of a previously published study^16,17^, a subset of the patients had been subjected to WES (n = 67) of tumor and blood samples and total RNAseq (n = 44) of tumor samples. Follow-up data have been updated since the first publication and ctDNA has been reevaluated using WGS (n = 119)^18^. Patients were categorized as metastatic if metastases were detected by computed tomography (CT) scan or other clinical follow-up after a cystectomy attempt (n = 31). Non-metastatic patients were disease-free with at least two years of follow-up (n = 79) after cystectomy. Nine patients had insufficient follow-up (< two years) or died within two years, and were excluded from all analyses comparing patients with and without metastatic disease. Three patients with non-successful cystectomy or death before the first CT scan were excluded from RFS curves. Pathological complete response was defined as T0,N0 after chemotherapy, while the non-invasive response was defined as T1,CIS,N0 or less. We performed TCRseq on blood samples taken at diagnosis (before administration of neoadjuvant chemotherapy). A subset of patients was included for TCRseq on tumor samples (n = 30) and longitudinal blood samples taken after chemotherapy, three weeks after cystectomy, and either when metastatic disease was detected or one year after cystectomy (n = 33). Biochemical laboratory measurements of blood cell counts were obtained through patient journals. The study’s second cohort is a subset of 30 patients from a larger study cohort of 156 patients diagnosed with NMIBC all receiving BCG immunotherapy. Blood samples collected before and after BCG along with treatment-naive tumor samples from 30 patients were subjected to TCRseq. Patients were categorized into early high-grade recurrence or progression (n = 15) if high-grade urothelial carcinoma was detected within two years after the end of BCG induction treatment or if patients progressed to MIBC at any time during follow-up. Patients with late or no high-grade recurrence (n = 15) were free of high-grade tumors for at least two years after the end of BCG induction treatment. Blood and tumor WES data were available for all patients in the full cohort (n = 156)^1^. For single-cell analysis, we included four patients undergoing cystectomy at the Department of Urology, Aarhus University Hospital, Denmark in May and June 2023. None of these patients were treated with chemotherapy prior to cystectomy.

### DNA and RNA extraction

DNA was purified from frozen buffy coat samples on QIAsymphony SP using QIAsymphony DSP DNA Midi Kit (QIAGEN). Tumor DNA was purified from fresh-frozen tumors or formalin-fixed paraffin-embedded tumors using Gentra Puregene Tissue Kit (QIAGEN) or AllPrep DNA/RNA Kit (QIAGEN), respectively.

Blood buffy coat RNA was purified from frozen buffy coat samples using miRNeasy Mini Kit (QIAGEN). Approximately 150 mm^3^ of frozen blood was cut out of a cryogenic tube using a scalpel and placed in a 2 mL Eppendorf tube. The cells were disrupted by mixing with 1.5 mL QIAzol Lysis Reagent before adding 140 µL chloroform. The remaining steps were performed according to protocol.

The quality of DNA and RNA was assessed using TapeStation5200 (Agilent) and the yield was determined using Qubit Fluorometric quantification (ThermoFisher Scientific) and DropSense96™ (Trinean) of the DNA and RNA, respectively.

### Library preparation and sequencing

AmpliSeq™ for Illumina ® TCR beta-SR panel was used to create TCR libraries for sequencing using 200 ng input DNA. The quality of TCR libraries was assessed using TapeStation4200 (Agilent) and Quibit Fluorometric quantification (ThermoFisher Scientific). The TCR libraries were paired-end sequenced on the Illumina NovaSeq6000 platform using SP and S1 flow cells (v1.5, 2x101 cycles), yielding an average of 42 million reads covering the CDR3β region (range 5-178 million).

Illumina Stranded Total RNA Prep with Ribo-Zero Plus kit was used to generate RNA libraries using 50 ng input RNA purified from buffy coat. The quality and yield of RNA libraries were assessed using TapeStation4200 (Agilent) and Quibit Fluorometric quantification (ThermoFisher Scientific). Sequencing was performed on the Illumina NovaSeq6000 platform using S2 flow cells (v1.5, 2x150bp), yielding an average of 305 million reads (range 101-357 million).

### TCRseq data analyses

The raw base call files were demultiplexed into FASTQ files using blcfastq (v2.20.0.422) from Illumina, allowing one mismatch in the index sequence. Subsequently, TCR clones were extracted from the data with MiXCR (v3.0.13)^39,40^ using the analyze amplicon function with 5-end v-primers, 3-end j-primers, and adapters present. Subsequently, clones with less than 50 reads were filtered out. These extremely low-frequency clones were characterized by abnormal length and frequent frameshifts, indicating that they were mainly sequencing artifacts. To have one TCR representing each clone, we removed non-productive TCR protein sequences from further analysis. Clones were defined based on nucleotide sequence unless otherwise mentioned. The normalized Shannon diversity index was calculated using the equation: Normalized Shannon diversity index = -1/log N *∑ (N, i = 1) p_i_ *log p_i_. Where N is the total amount of clones in a sample and p_i_ is the frequency of clone i. The fraction of expanded clones was quantified as the total frequency of clones above a frequency of 0.002. Overlapping TCR clones were determined based on the CDR3β nucleotide sequence. To reduce plot size, bubble plots of bulk blood TCR repertoires show top clones corresponding to 75% of the total repertoire frequency.

### RNAseq data analyses

Tumor RNAseq was previously quantified using Salmon^41^, utilizing Gencode annotations (v.33) on GRCh38 as in Lindskrog et al^42^. Differential gene expression analysis was performed in R using DESeq2 (v.1.38.3)^43^, which uses a Wald test. This analysis included 44 patients with MIBC, 21 with high TCR diversity, and 23 with low TCR diversity. Only protein-coding genes, excluding mitochondrial genes, were kept for analysis resulting in 18,665 genes from EnsDb.Hsapiens.v86 (v.2.99.0)^44^. Genes with a count of less than 10 in at least 21 samples (smallest group) were excluded. The top 2000 most variable genes were selected based on standard deviation. P-values were adjusted for multiple testing using the false discovery rate (FDR)^45^, and genes with an adjusted p-value below 0.25 were considered significant. Per category, the significantly upregulated genes (n = 153 for patients with low diversity, n = 39 for patients with high diversity) were used to calculate Reactome pathway enrichments using ReactomePA (v.1.42.0)^46^, which estimates enrichments based on a Hypergeometric model.

Blood RNAseq was quantified using Kallisto (v.0.48.0)^47^ utilizing Gencode annotations (v.37) on GRCh38. Transcript counts were collapsed into counts at the gene level (**Data S1**). Gene expression count data were normalized using EdgeR’s (v3.40.2)^48^ Trimmed-Mean of M-values. We downloaded the gene set from Travaglini et al.^30^ as a part of the C8 gene set collection from MSigDB and conducted a Gene Set Variation Analysis using GSVA (v.1.46.0)^49^. A GSVA score for naive and effector T cells was found by combining the “CD8 naive T cells” set with the “CD4 naive T cells” set, and the “CD8 effector memory T cells” set with the “CD4 effector memory T cells” set, respectively. We calculated the ratio of naive to effector T cells and compared patients with low and high TCR diversity.

### WES data analyses

Tumor and buffy coat WES data from 67 patients in the MIBC cohort, previously produced, were reanalyzed using the GRCh38 reference genome. Fastq files were trimmed using cutadapt and mapped with bwa-mem using the GRCh38 genome assembly. Duplicate reads were marked using MarkDuplicates from GATK and base quality scores were recalibrated (ApplyBQSR, GATK). Variants were called using Mutect2 and annotated using SnpEff (v4.3i). Finally, variants with a frequency below 5% (VAF < 5%), less than three alternate allele reads in the tumor, or a ROQ score below 30 (Phred-scaled probability that the variant alleles are not due to a read orientation artifact) were filtered out.

Additionally, 110 of the 156 patients in the NMIBC cohort had WES data captured with the Twist Human Core Exome Capture kit available. These samples were included for estimating T cell fractions. Reads were aligned and processed as described above. Of these 110 patients only 9 overlapped with the TCR analysis (total patients included = 131).

Based on germline WES data, we estimated blood T cell fractions using TcellExTRECT with default settings. For MIBC we used capture targets of TCRA genes from the SeqCapEZ MedExomeV1_hg19 capture kit. For NMIBC we estimated T cell fraction using the Twist human core exome capture targets.

For the tumor WES data, VCF-files were annotated using annovar (annotate_variation.pl, version from 2018-04-16)^50^, vcfR (v1.14.0)^51^, data.table (v1.14.8)^52^, and tidyverse (v2.0.0)^53^. Mutations were considered driver mutations if they were either 1) in a list of known driver mutations; 2) single nucleotide variants in tumor suppressor genes that were either predicted deleterious by MetaSVM^54^ or SIFT^55^, or annotated as being a stop-gain or splice mutation; 3) single nucleotide variants in oncogenes present at least three times in COSMIC (v90; cancer.sanger.ac.uk)^56^; or 4) any given mutation annotated as being either a non-synonymous, a stop-gain or a splice mutation that is present at least ten times in COSMIC. For the ten most mutated genes in MIBC from TCGA^21^, we counted the number of patients with and without driver mutations, with high and low TCR diversity, respectively. For each gene, we tested the difference in the number of driver mutations between patients with high and low TCR diversity using a Fisher’s Exact test.

CHIP was called based on WES data from buffy coat samples. Mutations were called using Mutect2 (GATK v4.4.0.0)^57^, using GnomAD^58^ variants as germline reference. A panel of normals was created from 25 tumor samples without CHIP mutations, found by running the pipeline on all tumor samples using a panel of normals generated from the 1000 genomes project^59^. Subsequently, variants were filtered using FilterMutectCalls, and variants with germline-like VAF were excluded (determined by non-significant deviance from VAF 0.5, binomial test). The remaining variants were annotated using VEP (v107)^60^. CHIP mutations were defined by a list of CHIP-specific mutations in 74 genes used to annotate CHIP by Bick et al.^23^. CHIP was called if a sample had at least one CHIP-related mutation with >2% VAF.

### WGS data analyses

Plasma WGS data from 119 patients with MIBC were used to detect ctDNA as described in Nordentoft et al.^18^. HLA type was called based on buffy coat WGS data from these 119 patients, using POLYSOLVER^61^.

To detect latent viruses we utilized plasma WGS data collected throughout treatment (total n = 973, median 8 samples per patient, median read count per patient = 950 million reads). We extracted unaligned reads from these samples and detected viral DNA using Kraken2^27^. A standard index created by the original authors was used as a reference (benlangmead.github.io/aws-indexes/k2). We used an index that contains DNA from other species to filter out reads originating from non-viral species, especially poor-quality human reads. EBV and CMV DNA detection was called for a patient if any of the patient’s samples contained more than 1 read identified as CMV or EBV with a distinct minimizer above 10. Fisher’s exact test was used to test for an association between the development of metastatic disease and CMV DNA detection.

### Annotation of TCR sequences

To annotate TCR clones we started by clustering baseline MIBC CDR3β amino acid sequences based on local and global sequence similarity using GLIPH2^62^ utilizing the provided TRB human v2.0 CD48 reference data and default parameters. WGS-based HLA type was included in the analysis. The analysis was independent of clone size, as clone frequency was excluded. TCR sequences of abnormal length were not included in the analysis (≧ 25 residues). High confidence clusters were established by selecting clusters with at least 3 unique sequences in at least 3 patients, and with significant Vβ gene enrichment bias, resulting in 12,831 clusters with a total of 63,299 CDR3β sequences. To infer an antigen target for the clusters, we reclustered the CDR3β sequences together with a dataset of high-confidence CDR3β sequences with known antigen targets obtained from VDJdb^63^. Only CDR3β-antigen pairs with a VDJdb confidence score above zero were included in the study, resulting in 4437 unique pairs. Clusters that were found in both runs were analyzed further. Note that this two-step process ensures that the CDR3β-antigen pairs do not bias cluster formation. Clusters sharing sequence motifs with a CDR3β-antigen pair were considered to target that antigen. Clusters with multiple targets were defined to target the most common named antigen. Fisher’s exact test was used to test for association between being included in a cluster with an inferred target and a clone being expanded, and a one-sided Fisher’s exact test was used to test for inferred target enrichment among expanded clones.

### Validation datasets

For TCGA we included all stage I-III patients with blood-derived normal samples (n = 262). The blood samples are chemotherapy-naive making them directly comparable to our baseline samples. Blood-derived normal WES bam files were downloaded from the GDC data portal and analyzed. Productive TCR beta sequences were extracted using MiXCR’s analyze shotgun functionality. For diversity estimation, patients with more than two reads and at least two distinct clones were included in the analysis. Below this threshold, repertoires will always be perfectly diverse (normalized Shannon diversity index = 1), and thus diversity can not be estimated. Blood T cell fractions were estimated using TcellExTRECT using exons from the corresponding capture kit (Agilent custom V2 Exome) with a median coverage threshold set to five.

For the IMvigor210 dataset, PBMC WES fastq files were acquired from EGA (EGAD00001004218). Reads were aligned and processed as described previously. Blood T cell fractions were estimated using TcellExTRECT using covered targets from Agilent SureSelect All Exon v5 (S04380110) overlapping TCRA genes. Productive TCR beta sequences were extracted using MiXCR’s analyze shotgun functionality. For diversity estimation, patients with more than two reads and at least two distinct clones were included in the analysis.

### Single-cell analyses

Fresh tumor and blood samples were collected from the Department of Urology, Aarhus University Hospital, Denmark. The samples were placed on ice and transferred to the laboratory for immediate processing. The tumor samples were washed with PBS and dissected using scalpels before transferring them into a gentleMACS C tube containing 3 mL PBS with 2% FBS and 1 mM EDTA. The samples were further dissociated using the gentleMACS m_intestine_01 program. Cells were filtered through a 70 µm and a 50 µm mesh and then centrifuged at 1300 rpm for 5 minutes at 4°C. The pellets were resuspended in 1 mL PBS with 2% FBS and 1 mM EDTA and subsequently filtered through 40 µm flowmi filters. T cells were isolated from the cell suspensions using the EasySep™ Human CD3 Positive Selection Kit II (STEMCELL™) according to protocol. T cells were isolated from 1mL of the blood samples using the EasySep™ Direct Human T Cell Isolation Kit (STEMCELL™) according to protocol. The concentration and viability of the samples were measured using Via1-Cassettes™ on a Nucleocounter® NP-3000™. Samples with a concentration below 700,000 cells/mL were centrifuged for 5 min at 250G at RT and resuspended in PBS to a concentration of 1,000,000 cells/mL. Samples with a concentration above 1,200,000 were diluted using PBS to a concentration of 1,000,000 cells/mL.

Chromium Next GEM Single Cell 5’ Reagent Kits v2 (Dual Index) from 10X Genomics was used for single cell sequencing and quality was measured under way using Tapestation HS-D5000. The quantity of the final libraries was measured using Qubit. All libraries were paired-end sequenced on the Illumina NovaSeq 6000 platform using SP and S1 flow cells (v1.5, 300 cycles).

The datasets were pre-processed (demultiplexed, reads aligned and filtered, barcodes and UMIs counted, and TCR clones determined) using Cell Ranger (10X Genomics). The data were filtered based on unique features and mapping to mitochondria genes, keeping cells with >200 and <2500 unique features and <10% mitochondria mapping using Seurat (v4.4.0)^64^. Clones with a TCR β sequence were included for analyses. The cells were annotated using CellTypist^29^ with the model Immune All Low which has high resolution and includes 98 different immune cell types. The cell type of a T cell clone was called using majority vote and only cells annotated as T cells were included in the analyses.

### Statistical analysis

All pairwise comparisons are tested using a Wilcoxon Rank Sum test, except when comparing paired data, then a Wilcoxon Signed Rank test was used. For time-series laboratory count data, unnormalized values were used. For boxplots, the center line represents the median, box limits represent upper and lower quartiles, and whiskers represent 1.5 times the interquartile range. For survival analyses, differences between Kaplan-Meier curves are tested using a likelihood ratio test, and the HR and 95% confidence interval are calculated using Cox proportional hazard regression. Cox proportional hazard regression was used for multivariate analyses. Correlations were tested using Spearman’s rank correlation. If other tests were used they are described in the relevant method section. All statistical tests were two-sided unless otherwise stated. When relevant multiple testing correction was performed using FDR. P-values or FDR-adjusted p-values were considered significant when below 0.05 unless stated otherwise.

## Data availability

Processed data produced for this publication, including TCRseq, RNAseq, and single-cell RNAseq, and summary data to create all figures are available as **Data S1-9** and **Table S2**. The raw sequencing data generated in this study are not publicly available as this compromises patient consent and ethics regulations in Denmark.

## Code availability

All code used to produce figures is available at the project github: https://github.com/nbirkbak/Bladder-TCRseq.

## Acknowledgments

All of the computing for this project was performed on the GenomeDK cluster. We thank GenomeDK and Aarhus University for providing computational resources and support that contributed to these research results. The authors acknowledge funding received in support of the project from AstraZeneca, The Danish Cancer Society, The Novo Nordisk Foundation, the Lundbeck Foundation, and the Aarhus University Research Foundation.

## Author contributions

A.K., N.K., and R.I.J. contributed equally. N.J.B. and L.D. conceived the study design. A.K., N.K., R.I.J., I.N., and D.R. performed data integration. A.K., N.K., R.I.J., and J.A. performed statistical analyses. I.N. coordinated and supervised bulk TCRseq. N.K. performed single-cell sequencing and RNAseq on blood samples. K.B.-D. and T.S. selected samples for bulk TCRseq and collected clinical follow-up information. J.B.J. provided samples for single-cell sequencing. A.K., N.K., R.I.J., N.J.B., and L.D. drafted the manuscript. N.J.B. and L.D. supervised the study. All authors provided feedback and interpretation of results, and all authors approved the final version of the manuscript.

## Supplemental information titles and legends

Document S1: Figures S1-S12 and Table S1.

Table S2: Excel file containing the source data reproduce analyses and create all figures.

Data S1: Summarized gene expression counts as outputted by Kallisto for the RNAseq of blood samples.

Data S2-S9: Single-cell RNAseq counts from blood and tumor for four patients (two files per patient, one for blood and one for tumor).

