## Supplementary material for "Low T cell diversity is associated with poor outcome in bladder cancer: a comprehensive longitudinal analysis of the T cell receptor repertoire": Document S1

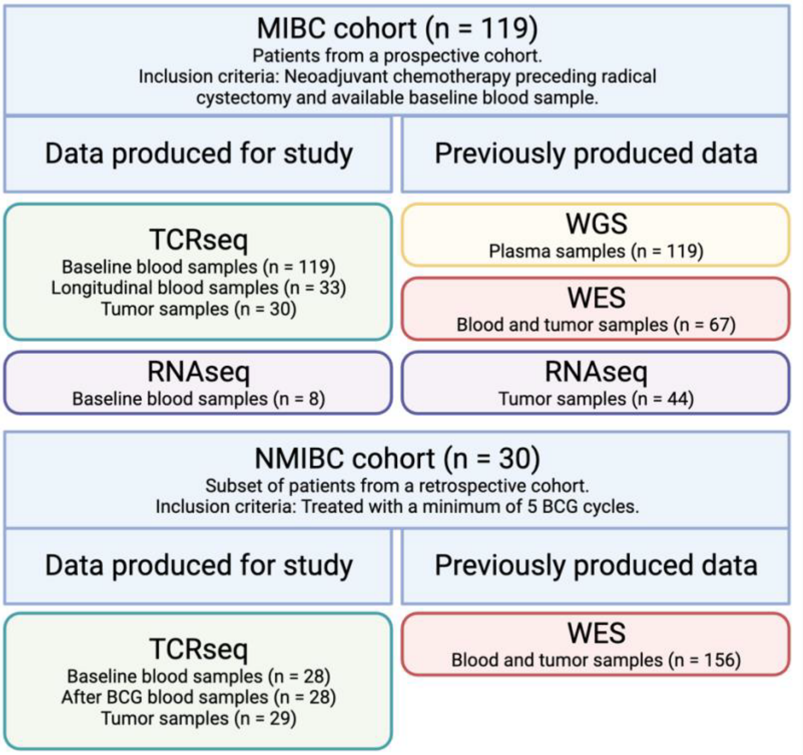


**Figure S1 | Sample overview**, related to **Figure 1**. Overview of analyzed samples for both cohorts. Created with BioRender.com.

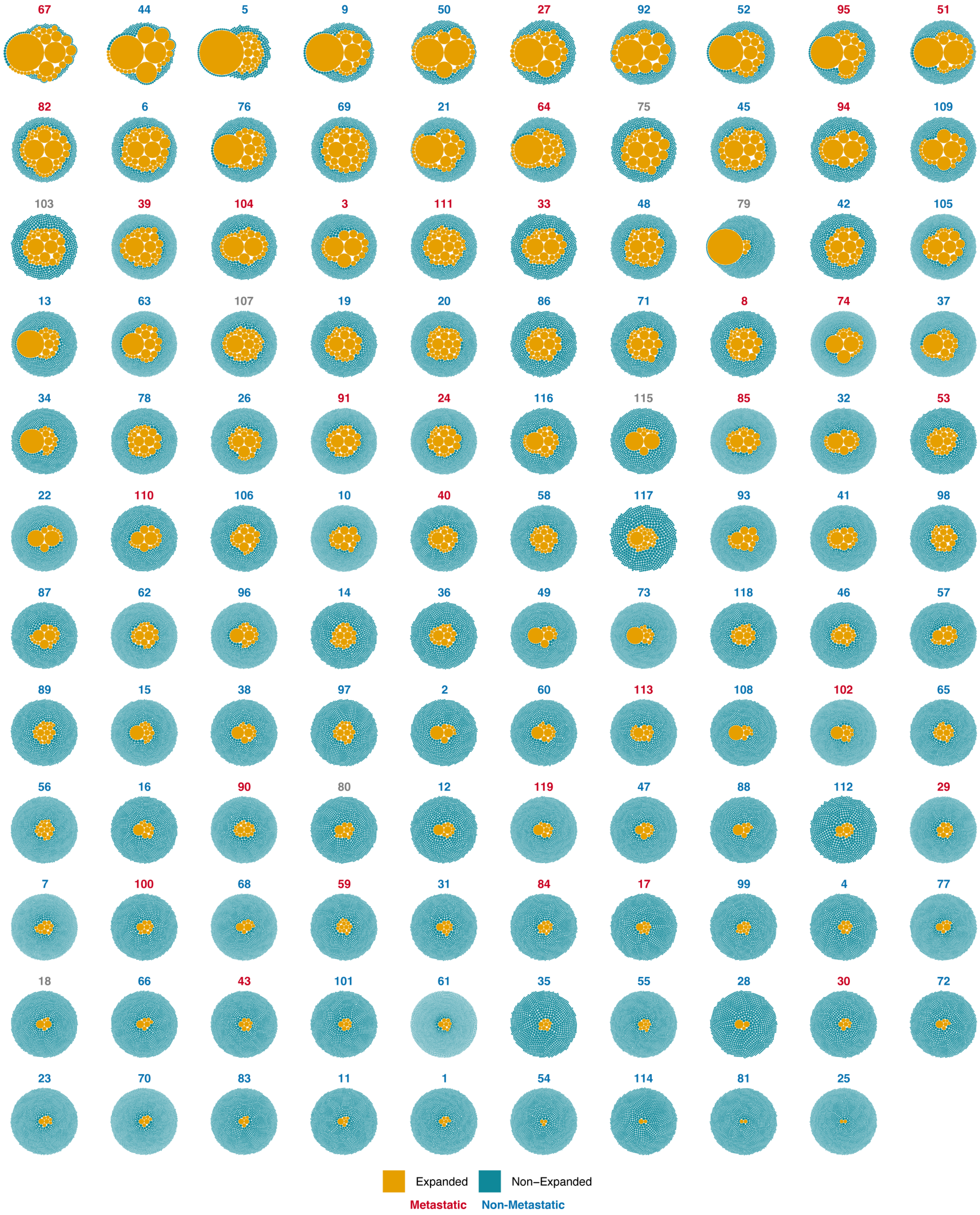


**Figure S2 | Bubble plots of baseline blood samples from the MIBC cohort**, related to **Figure 1**. One plot per patient. Each bubble represents a single TCR clone, with the size of the bubble representing the clone size. Colored by the size threshold for expansion. Patient labels colored by later development of metastatic disease.

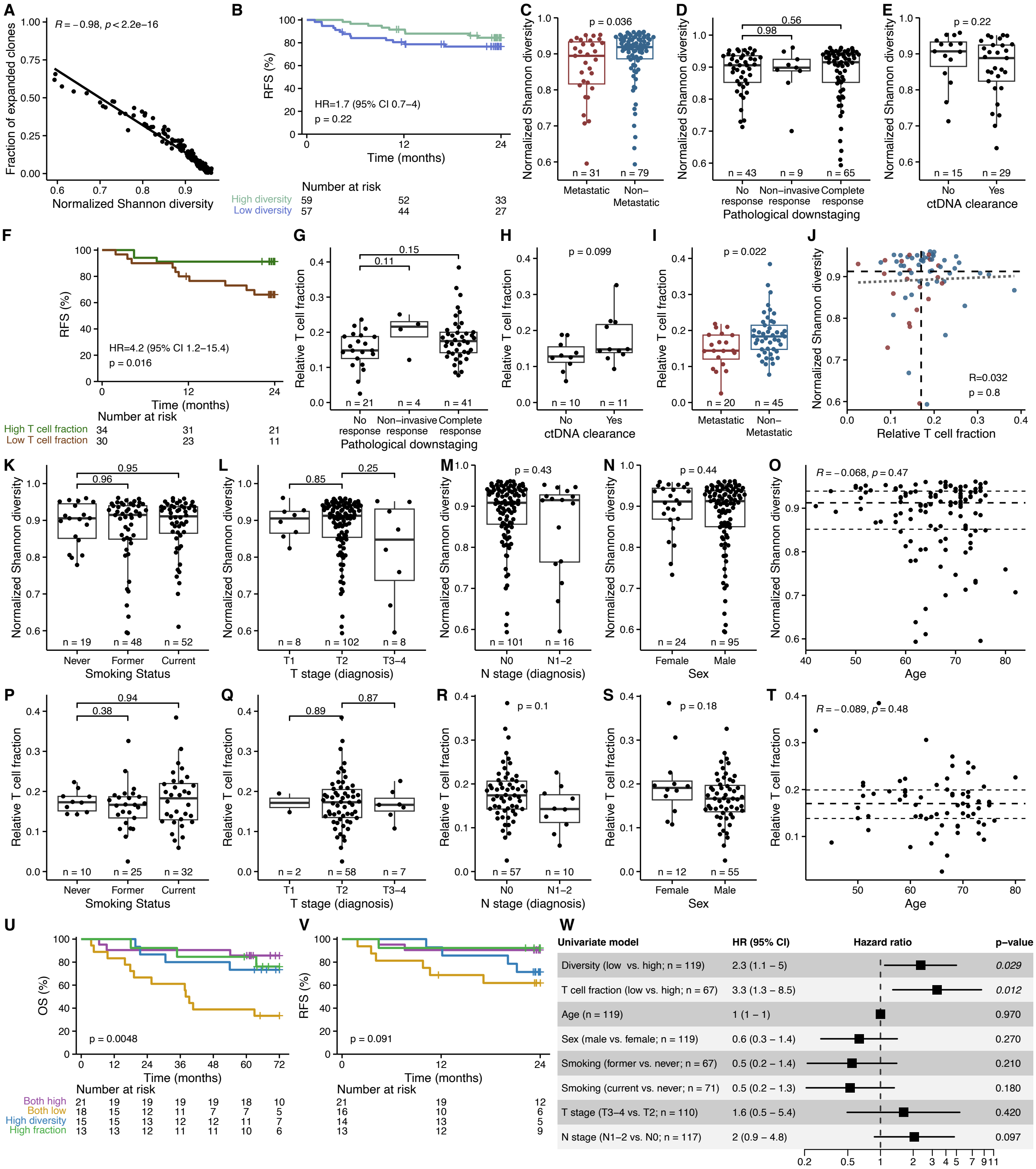

**Figure S3 | Baseline peripheral TCR features, relative T cell fraction and clinical parameters in MIBC**, related to **Figure 2**. **A,** Scatter plot showing a negative correlation between the fraction of expanded clones and normalized Shannon diversity index. **B,** Survival analysis associating TCR diversity (median split) with RFS. **C,** Box-plot comparing normalized Shannon diversity index between patients who develop metastatic disease during follow-up to those who do not. **D,** Association between normalized Shannon diversity index and pathological downstageing. **E,** Association between normalized Shannon diversity index and ctDNA clearance. **F,** Survival analysis associating relative T cell fraction (median split) with RFS. **G,** Association between relative T cell fraction and pathological downstageing. **H,** Association between relative T cell fraction and ctDNA clearance. **I,** Box-plot comparing relative T cell fraction between patients who develop metastatic disease during follow-up to those who do not. **J,** Scatter plot showing the Spearman correlation between normalized Shannon diversity index and relative T cell fraction indicated by the gray dotted line. Dashed lines indicate the medians. **K-O,** Association between baseline normalized Shannon diversity index and smoking status (**K**), T stage (**L**), N stage (**M**), sex (**N**), and age (**O**, dashed lines indicate quartiles). **P-T,** Association between relative T cell fraction and smoking status (**P**), T stage (**Q**), N stage (**R**), sex (**S**), and age (**T**, dashed lines indicate quartiles). **U-V**, Survival analyses of normalized Shannon diversity index combined with relative T cell fraction, both split by median. High diversity indicates high normalized Shannon diversity index and low T cell fraction. High fraction indicates high T cell fraction and low normalized Shannon diversity index. Survival analysis using OS (**U**) and RFS (**V**). **W,** Forest plot showing univariate models of TCR diversity, T cell fraction, age, sex, smoking status, T stage, and N stage. CI: confidence interval.

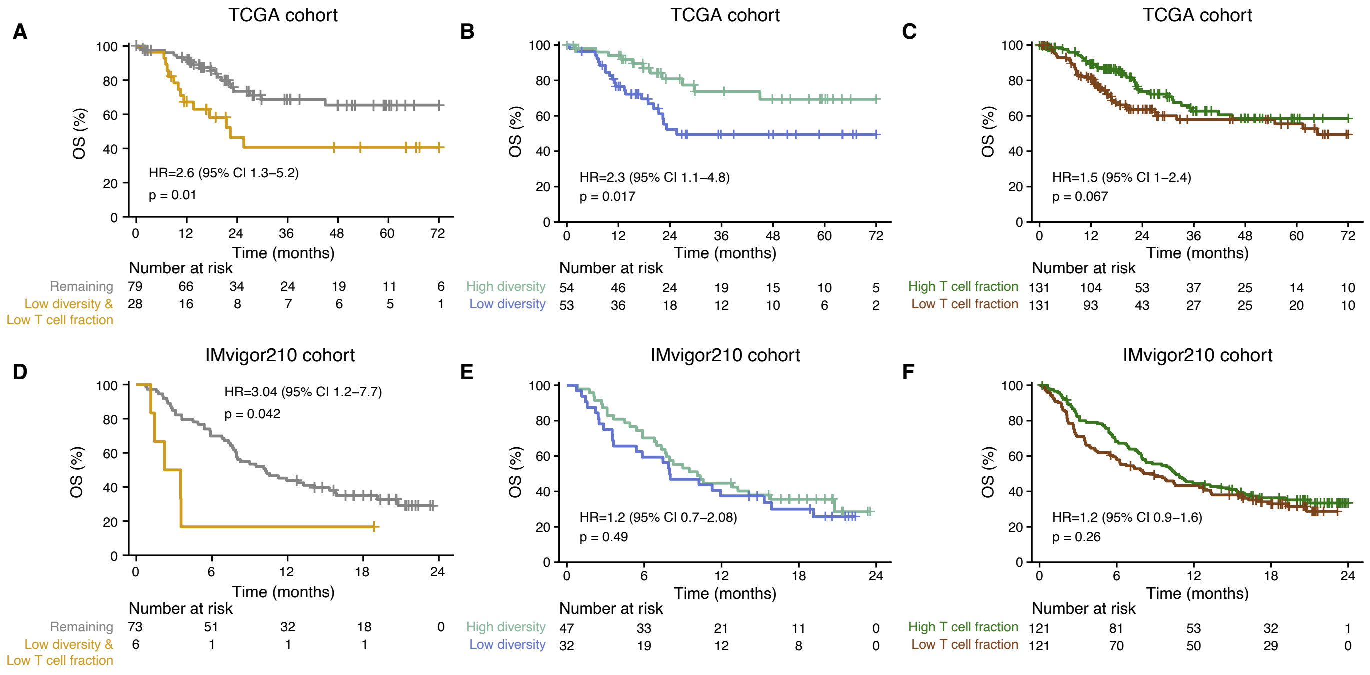


**Figure S4 | Analysis of TCR diversity and T cell fraction in independent cohorts**, related to **Figure 2**. **A-C,** Stage I-III MIBC from TCGA (n = 262). Normalized Shannon diversity index was determined by extracting TCR-beta sequences from germline WES samples. Only 107 patients had sufficient coverage of the CDR3 region to determine diversity. Survival analysis comparing OS for patients with low TCR diversity (median split) combined with low T cell fraction (median split) against the remaining patients (**A**), low and high (median split) TCR diversity (**B**), and low and high (median split) T cell fraction (**C**). **D-F,** Patients with metastatic bladder cancer from IMvigor210 (n = 242, n diversity = 79). Survival analysis comparing OS for patients with low TCR diversity (median split) combined with low T cell fraction (median split) against the remaining patients (**D**), low and high (median split) TCR diversity (**E**), and low and high (median split) T cell fraction (**F**). CI: confidence interval.

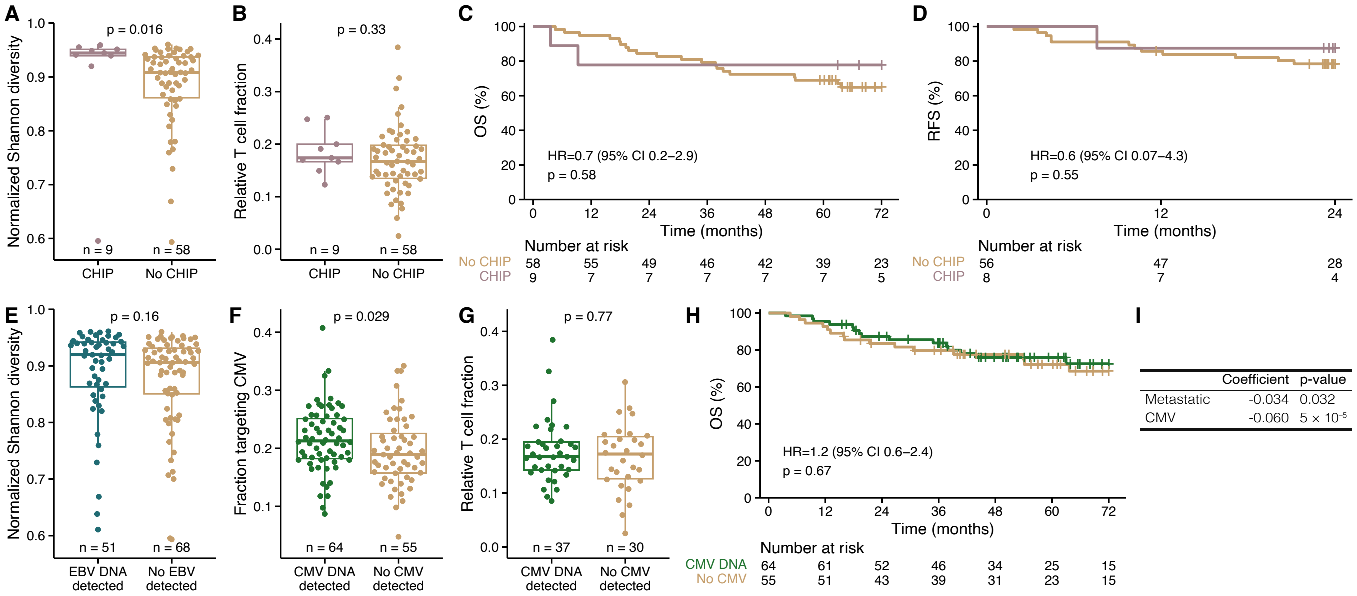


**Figure S5 | Analyses of CHIP and latent viral infections**, related to **Figure 3**. **A-D,** Analysis of CHIP in patients with MIBC. CHIP was called based on the detection of somatic, CHIP-associated mutations found in germline WES data. **A,** Box-plot showing that patients with detectable CHIP have higher TCR diversity. **B,** Box-plot showing no difference in the relative T cell fraction between patients with and without detectable CHIP. **C-D,** Survival analyses showing that detectable CHIP does not affect OS (**C**) or RFS (**D**). **E-H,** Detection of CMV and EBV DNA in patients with MIBC. CMV and EBV DNA were detected based on WGS data from plasma samples using Kraken2. **E,** Box-plot showing no difference in the normalized Shannon diversity index between patients with and without detectable EBV DNA in plasma. **F,** Box-plot showing that patients with detectable CMV DNA have a higher fraction of TCR clones targeting CMV antigens. Quantified as the amount of clones targeting CMV out of all clones with an inferred target. **G,** Box-plot showing no difference in the relative T cell fraction between patients with and without detectable CMV DNA. **H,** Survival analysis showing that latent CMV infection was not associated with OS. **I,** Coefficients and p-values for a linear regression model to predict normalized Shannon diversity index based on metastatic disease and CMV detection (MIBC, n = 119). CI: confidence interval.

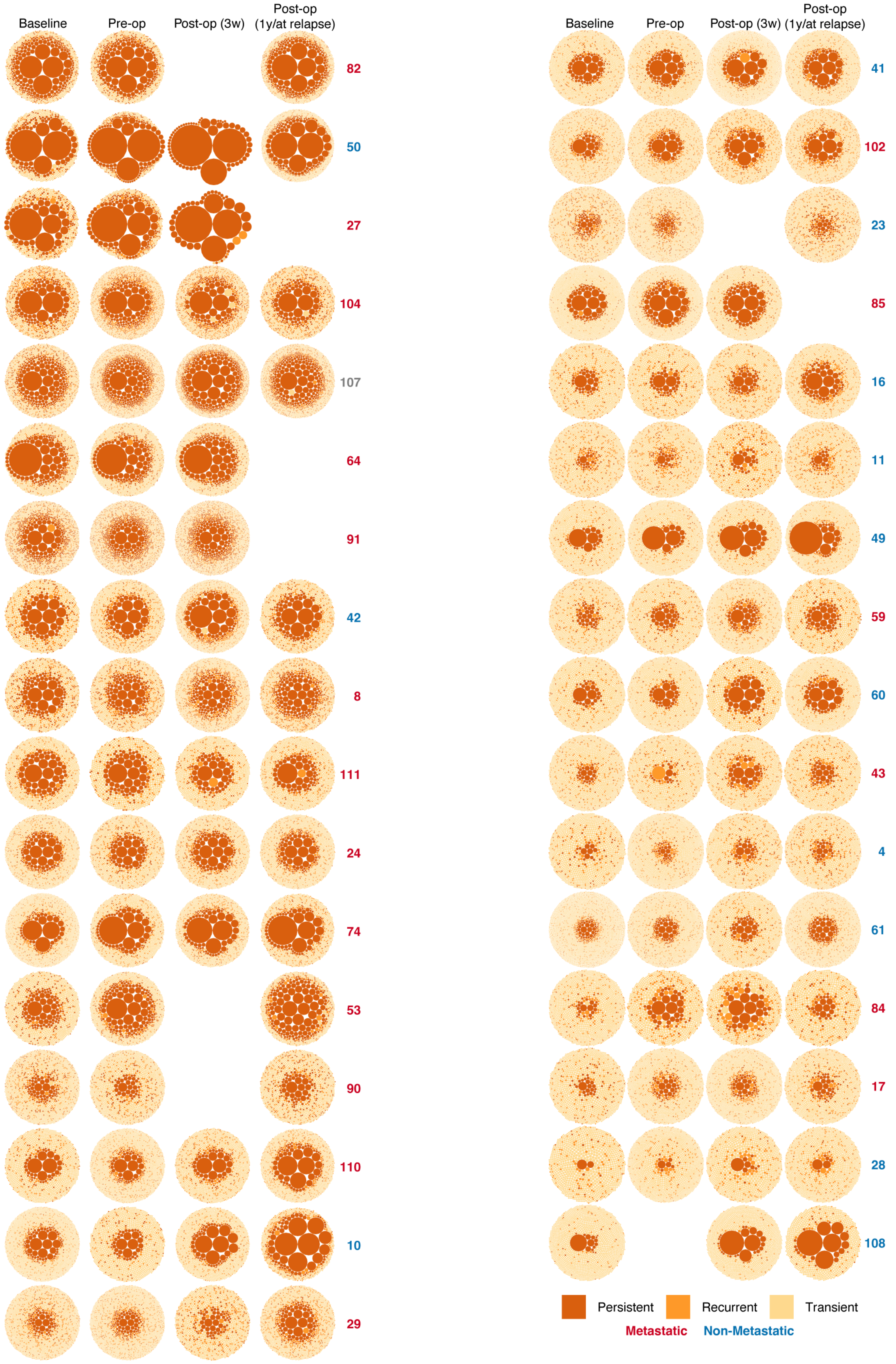


**Figure S6 | Bubble plots for longitudinal blood samples from the MIBC cohort**, related to **Figure 4**. One plot per patient per available time point. Each bubble represents a single TCR clone, with the size representing the clone size. Colored by clones being persistent, recurrent, or transient. Patient labels colored by later development of metastatic disease.

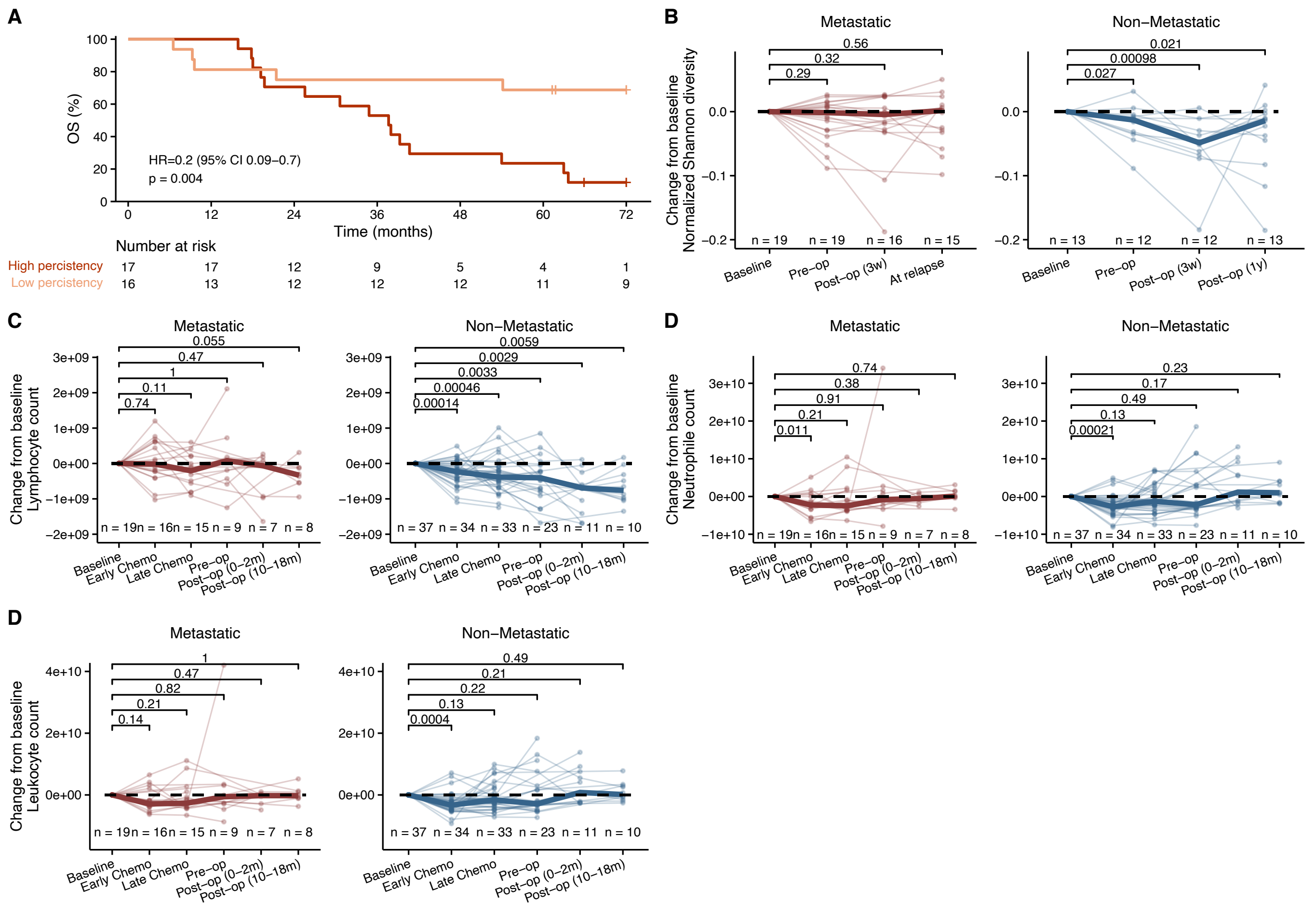


**Figure S7 | Analysis of persistency and blood lab measurements in MIBC**, related to **Figure 4**. **A,** Survival analysis showing that high persistency (above median amounts of persistent and recurrent clones) is associated with OS. **B-E,** Change in immune landscape throughout treatment for patients with and without metastatic disease separately; normalized Shannon diversity index (**B**), lymphocyte counts (**C**), neutrophil counts (**D**), and leukocyte counts (**E**). Early chemo represents samples taken within one month of initiation, and late chemo represents the remaining samples. When multiple samples were available the average count was used. The thick lines indicate the median. Tests are based on the raw diversity measures or lab counts.

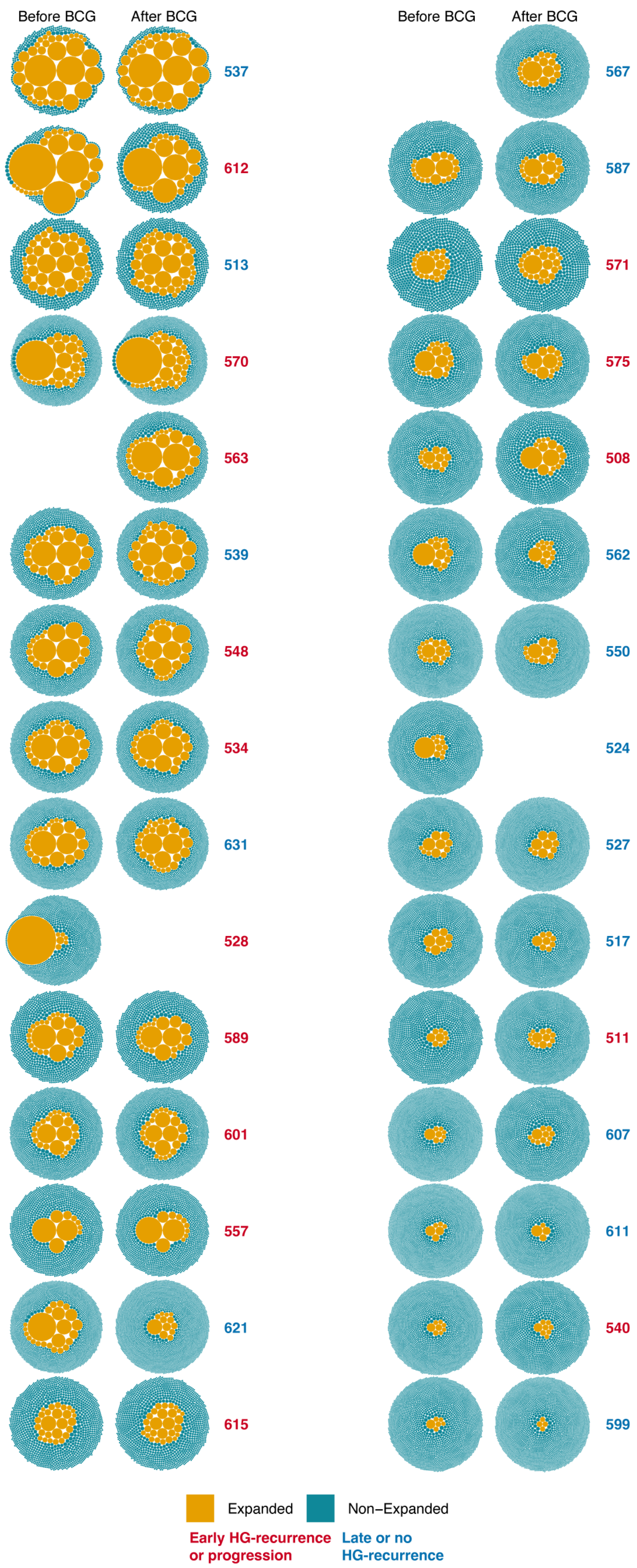


**Figure S8 | Bubble plots of blood samples from the NMIBC cohort.** One plot per patient per available time point. Each bubble represents a single TCR clone, with the size of the bubble representing the clone size. Colored by the size threshold for expansion. Patient labels colored by early HG recurrence or progression, or late or no HG recurrence. HG: high-grade.

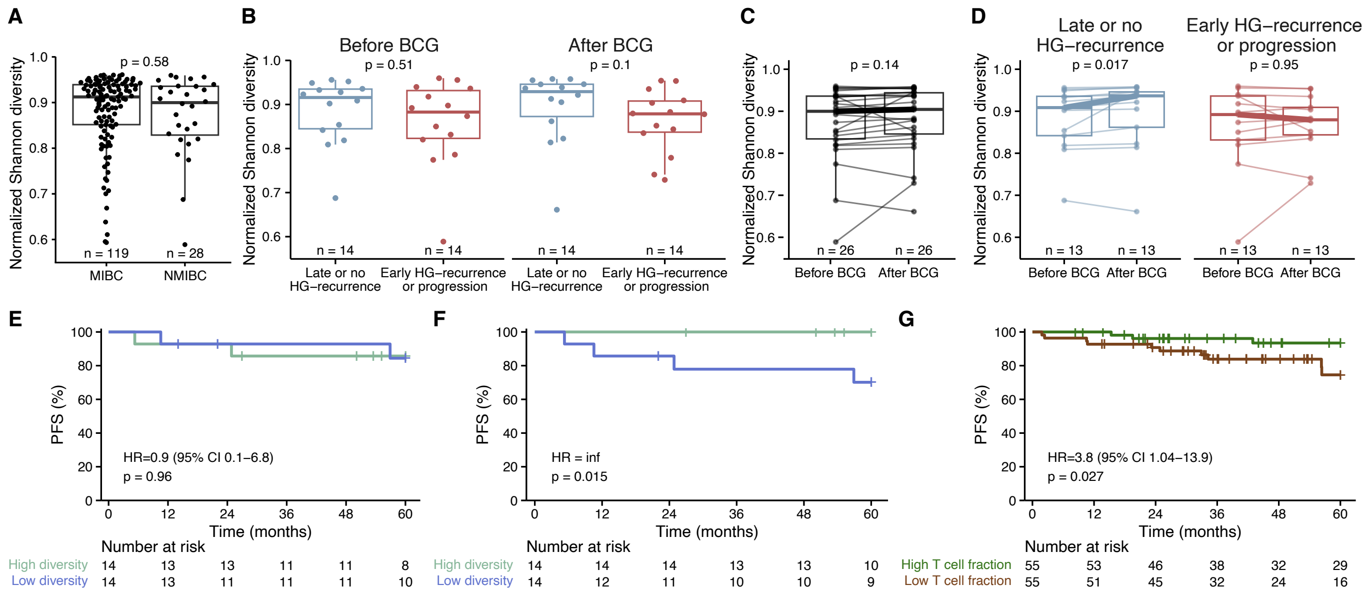


**Figure S9 | TCR repertoires in NMIBC. A,** Box-plot comparing normalized Shannon diversity index between patients with MIBC and NMIBC at baseline. **B-F,** Investigation of TCR diversity in patients with NMIBC (n = 30), treated with BCG immunotherapy. **B,** Box-plot comparing normalized Shannon diversity index before and after BCG, between patients with late or no HG recurrence and patients with early HG recurrence or progression at any time. **C,** Box-plot showing the paired change in normalized Shannon diversity index for each patient before and after BCG. **D,** Box-plot showing the change in normalized Shannon diversity index for each patient before and after BCG, split by patients with late or no HG recurrence and patients with early HG recurrence or progression. **E-F,** Survival analyses investigating the association of PFS and TCR diversity (median split) before (**E**) and after (**F**) BCG. **G,** Survival analysis investigating the association of PFS and relative T cell fraction (median split). T cell fraction was estimated on WES data from an extended cohort of patients with NMIBC (n = 110). HG: high-grade. CI: confidence interval.

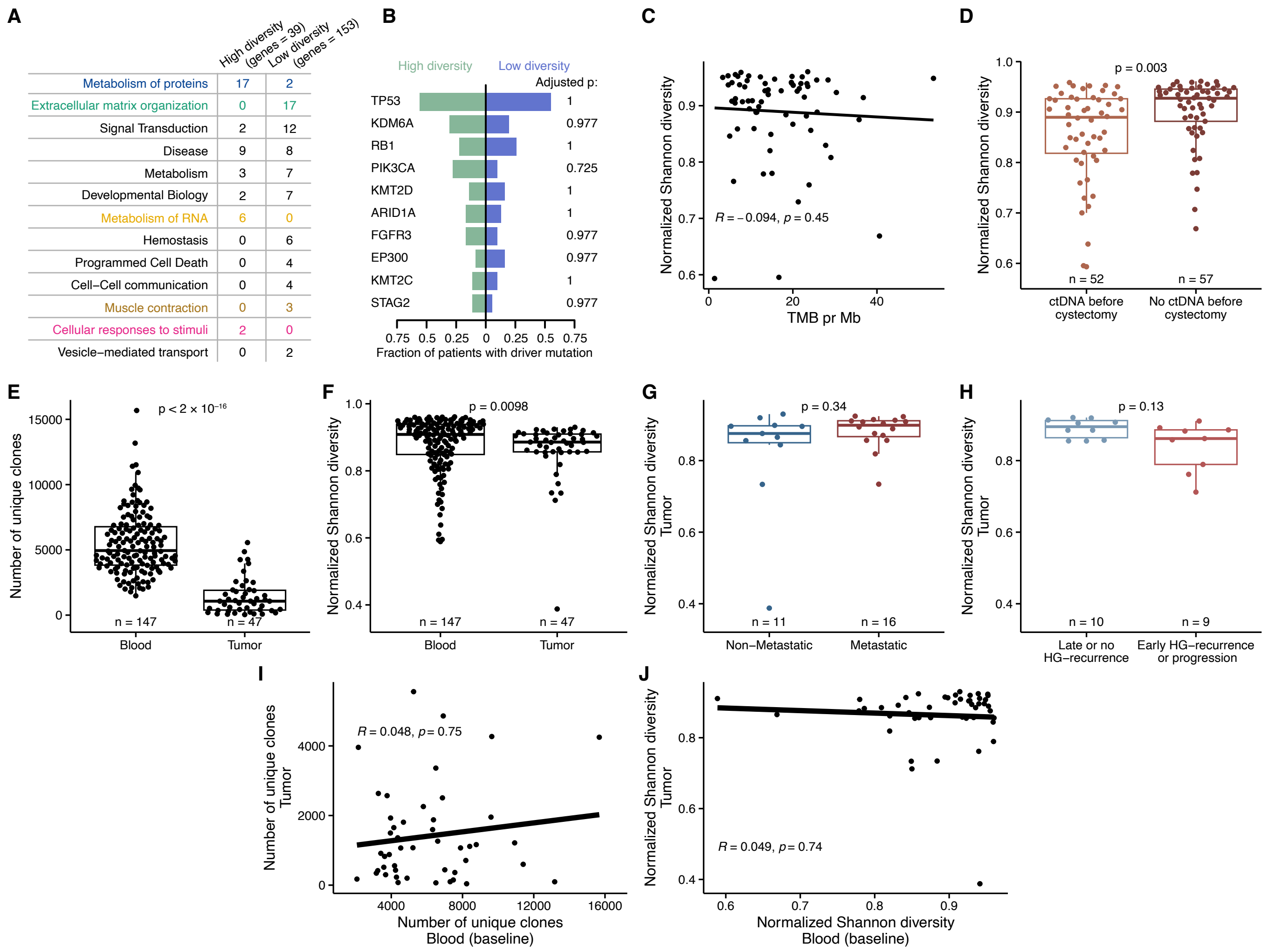


**Figure S10 | Association of peripheral TCR repertoires and tumor biology or tumor TCR repertoires**, related to **Figure 5**. **A-D,** Analysis of peripheral TCR diversity relative to tumor biology. **A,** Summary of significant pathways from the Reactome pathway analysis in **Figure 5B**. **B,** Tree-plot showing the difference in the number of driver mutations between patients with high and low TCR diversity in ten bladder cancer driver genes. **C,** Scatter plot showing the correlation between normalized Shannon diversity index and TMB. **D,** Box-plot comparing the normalized Shannon diversity index between patients with and without detectable levels of ctDNA before cystectomy. **E-J,** Joint analysis of paired tumor and baseline repertoires for patients with MIBC (blood n = 119, tumor n = 28) and NMIBC (blood n = 28, tumor n = 19). **E,** Box-plot comparing the number of unique clones in blood and tumor. **F,** Box-plot comparing the normalized Shannon diversity index in blood and tumor. **G,** Box-plot comparing normalized Shannon diversity index between tumors from patients with MIBC, with and without metastasis. **H,** Box-plot comparing normalized Shannon diversity index between tumors from patients with NMIBC, with late or no HG recurrence and early HG recurrence or progression. **I,** Scatter plot showing the correlation between the number of unique clones in the tumor and in the blood. **J,** Scatter plot showing the correlation between the normalized Shannon diversity index in the tumor and in the blood. HG: high-grade. TMB: tumor mutation burden.

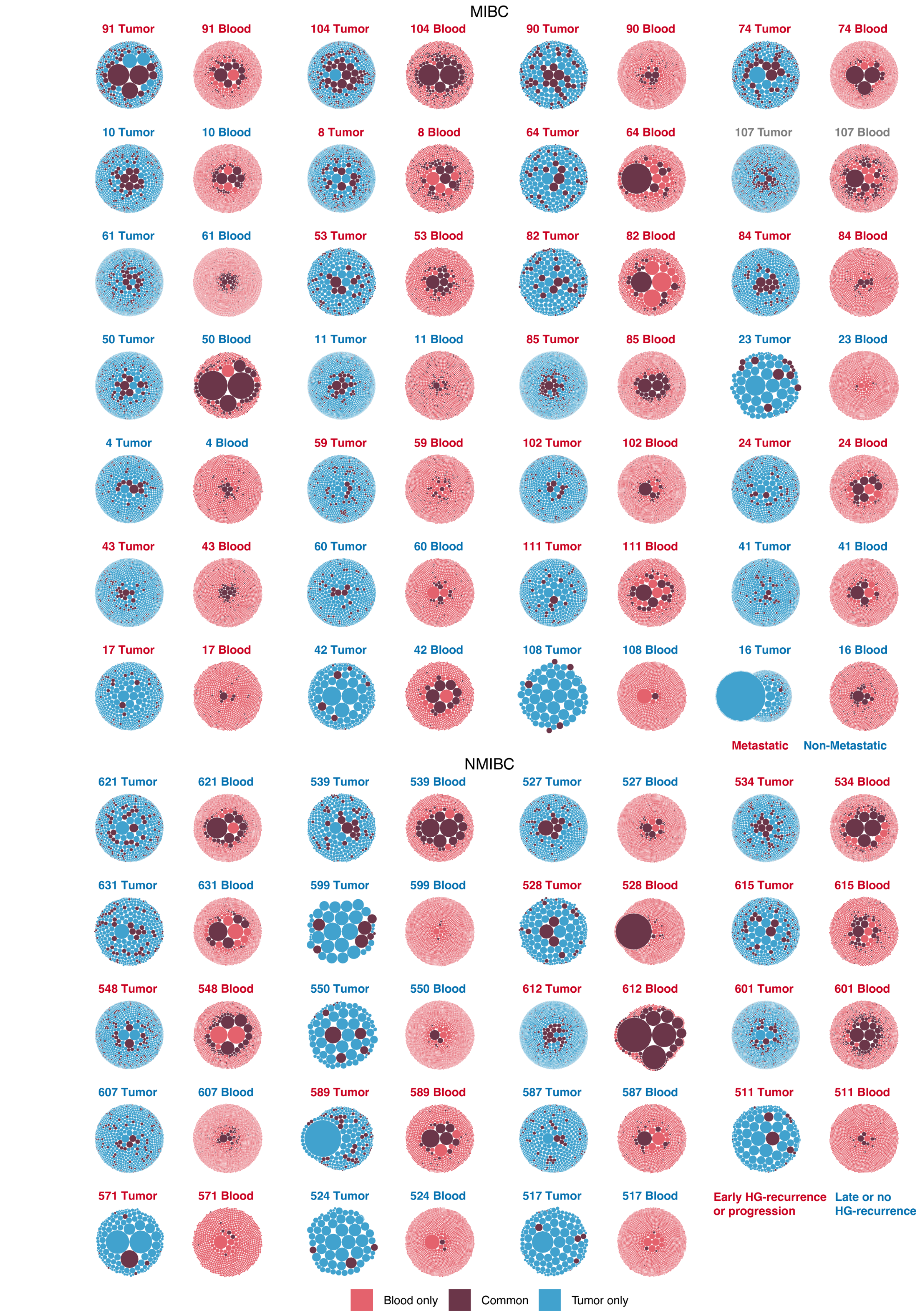


**Figure S11 | Bubble plots for tumor and blood samples from the MIBC and NMIBC cohorts**, related to **Figure 5**. Two plots per patient (tumor and baseline blood). Each bubble represents a single TCR clone, with the size representing the clone size. Colored by the clones being either unique to the tumor or blood or found in both (common). Patient labels colored by later development of metastatic disease (MICB cohort), or by early HG recurrence or progression, or late or no HG recurrence (NMIBC cohort). HG: high-grade.

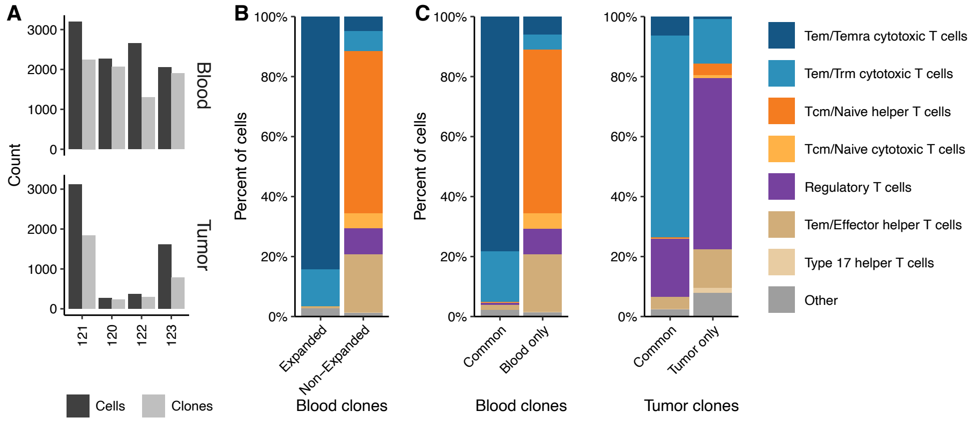


**Figure S12 | Single-cell analyses**, related to **Figure 6**. **A,** Bar plots showing the number of T cells and unique clones for blood and tumor. **B,** Bar plots showing the distribution of cell types split by expanded and non-expanded clones. **C,** Bar plots showing the distribution of cell types in blood and tumor clones, respectively, for both common and unique clones. Other includes: Trm cytotoxic T cells, Memory CD+ cytotoxic T cells, Treg(diff), Follicular helper T cells, Type 1 helper T cells, CRTAM+ gamma-delta T cells, gamma-delta T cells, MAIT cells, Cycling T cells, CD8a/b(entry), and Double-positive thymocytes. Tem: effector memory T cell. Temra: effector memory T cell reexpressing CD45RA. Trm: tissue-resident memory T cell. Tcm: central memory T cell. Treg(diff): differentiating regulatory T cell. MAIT: mucosal-associated invariant T cell. CD8a/b(entry): developing CD8 alpha-beta T cell (late double-positive stage).

**Table S1 | Summary of patient characteristics**, related to **Figure 1**

| | **MIBC (n = 119)** | **NMIBC (n = 30)** |
|---|---|---|
| **Sex** | | |
| Female | 24 (20.2%) | 7 (23.3%) |
| Male | 95 (79.8%) | 23 (76.7%) |
| **Age*** | | |
| Mean (SD) | 70 (8) | 70 (10) |
| Median [min, max] | 70 [40, 80] | 70 [50, 80] |
| **Smoking status** | | |
| Current | 52 (43.7%) | 17 (56.6%) |
| Former | 48 (40.3%) | 11 (36.7%) |
| Never | 19 (16.0%) | 2 (6.7%) |
| **T stage at diagnosis** | | |
| Ta | 0 (0%) | 18 (60.0%) |
| T1 | 8 (6.7%) | 12 (40.0%) |
| T2 | 102 (85.8%) | 0 (0%) |
| T3 | 1 (0.8%) | 0 (0%) |
| T4a | 4 (3.4%) | 0 (0%) |
| T4b | 3 (2.5%) | 0 (0%) |
| Tx | 1 (0.8%) | 0 (0%) |
| **N stage at diagnosis** | | |
| N0 | 101 (84.9%) | 30 (100%) |
| N1 | 13 (10.9%) | 0 (0%) |
| N2 | 3 (2.5%) | 0 (0%) |
| Missing | 2 (1.7%) | 0 (0%) |
| **Recurrence**** | | |
| Recurrence | 31 (26.0%) | 15 (50.0%) |
| No recurrence | 79 (66.4%) | 15 (50.0%) |
| Follow-up < two years | 9 (7.6%) | 0 (0%) |

* MIBC: age at diagnosis; NMIBC: age at BCG induction
** MIBC: detection of metastasis; NMIBC: early high-grade recurrence (< two years) or progression
SD: standard deviation
